# On the formation of lipid nanoparticles and encapsulation of RNA: kinetics, dynamics, and self-assembly regimes

**DOI:** 10.64898/2026.11.14.766710

**Authors:** Anas Aljabbari, Heather Binion, Harsa Mitra, Sophia R. Dasaro, Lucas D. Johnson, Titania Bethiana, Caleb F. Fretz, Dongjie Jia, Nicholas Zolnierczuk, Gabriel Harris, Mojhdeh Baghbanbashi, Andres Barrio-Zhang, David Perez Hererra, Ann Terry, Xiaobing Zuo, Brian K. Wilson, Arezoo Ardekani, Kurt Ristroph

## Abstract

RNA-containing lipid nanoparticles (LNPs) form through self-assembly triggered by micromixing of an organic lipid solution against an aqueous antisolvent, leading to supersaturation and consequent nanoprecipitation. The kinetics of this process remain poorly understood. Here, we employ millisecond-resolved flowthrough small-angle X-ray scattering (SAXS) coupled to a multi-inlet vortex mixer (MIVM), confined impinging jets mixer (CIJM) and a T-junction mixer, to experimentally investigate the earliest stages of LNP self-assembly. Ordered LNPs show a characteristic high-*q* feature indicative of the ionizable lipid-RNA complex; changes in the evolution, shape, and intensity of this peak and the overall particle size in the low-*q* feature reveal changes in particle assembly and RNA cargo encapsulation. Results captured within milliseconds of mixing show that under turbulent flow (water/ethanol ratio 3:1), LNPs are fully formed by as early as 12 ms, including liquid-crystalline internal organization that indicates RNA encapsulation. Adjusting the water/ethanol ratio to 1:1 results in LNPs with a less ordered nanostructure, in line with reports that downstream processing plays a major role in the final structure of particles formed under these solvent conditions. These findings shed light on the mechanisms by which mixing conditions influence LNP formation. We propose a framework of four distinct self-assembly pathways defined by mixing time (i.e. faster or slower than the O(10ms) self-assembly time) and loading strategy (i.e. RNA introduced during the LNP self-assembly step, or in a second step). This provides a basis for understanding how mixing and loading strategies govern LNP formation and homogeneous RNA encapsulation, particularly for high molecular weight RNA payloads. The results establish process-structure-function relationships that support rational optimization for LNP manufacturing processes.

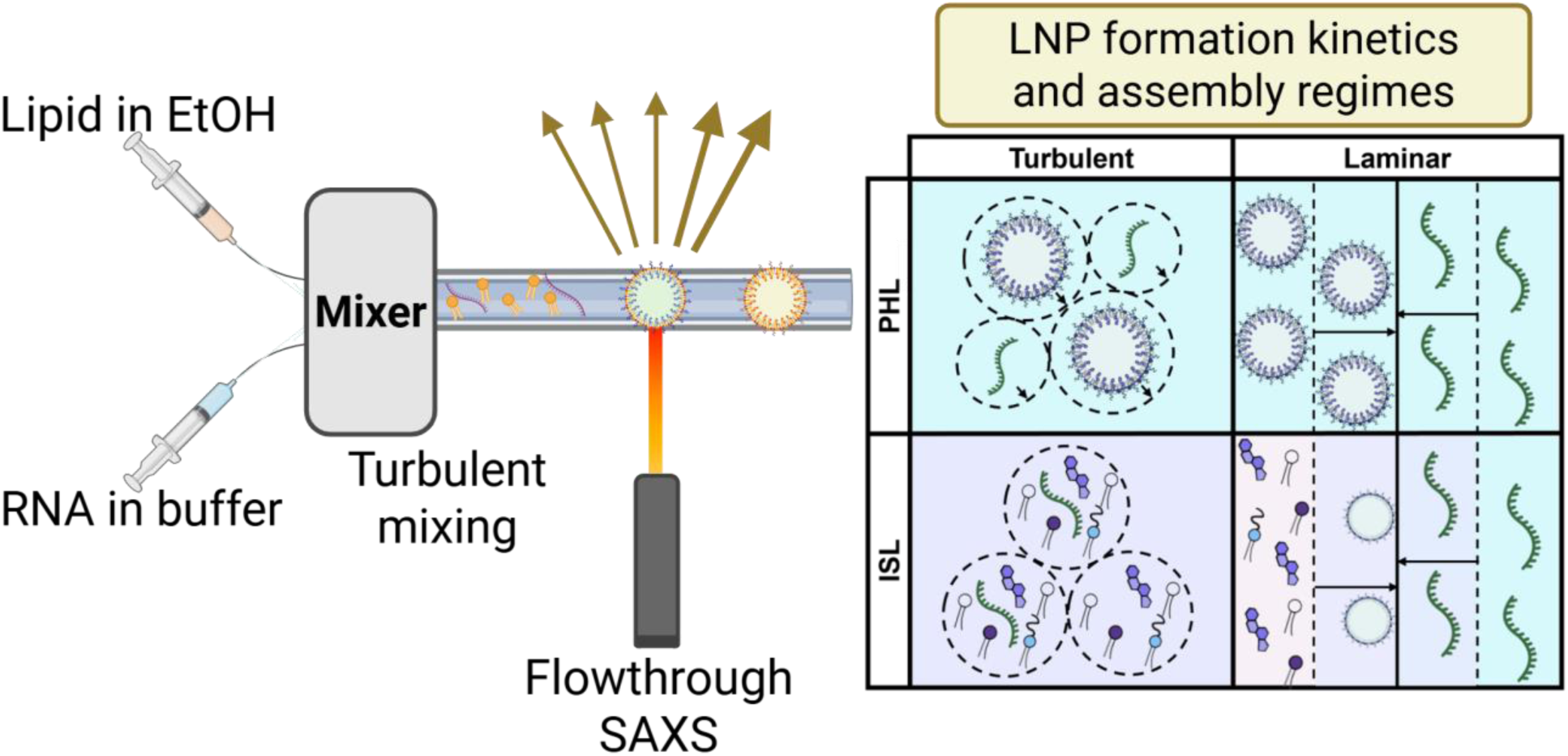

---

Lipid nanoparticles (LNPs) have emerged as a versatile delivery vehicle for nucleic acid therapeutics such as siRNA and mRNA that enable both protection of RNA cargo and efficient cellular transfection^1–3^. The biological performance of LNPs is strongly influenced by physicochemical and structural properties such size, polydispersity index (PDI), surface charge, internal liquid crystalline (LC) structure and overall particle morphology. Maintaining control over these critical quality attributes is therefore a central concern to LNP formulation and process design. While computational approaches, including molecular dynamics simulations and population balance equation models, have provided valuable insight into LNP self-assembly, experimental interrogation of the underlying kinetics remains limited due to the challenges associated with measuring self-assembly processes occurring on millisecond timescales ^4–6^. A recent report noted that experimentally resolving the kinetics of LNP self-assembly “would require millisecond-scale measurements, which is challenging.”^5^

We here present millisecond-scale measurements of LNP self-assembly kinetics, using flowthrough small-angle X-ray scattering (SAXS), and show that LNP formation concludes within ∼10 milliseconds post-mixing in turbulent mixers. We further demonstrate how variations in formulation and process parameters lead to subtle, but significant, differences in the regimes of self-assembly. By resolving the timescales required for LNP formation post-mixing, these findings can enable precise design and timing of downstream processing steps as well as establishing a rational basis for more advanced formulation engineering, such as homogeneous distribution of high molecular weight cargo or the co-encapsulation of multiple RNAs.

Clinical LNP formulations have four lipid components, listed in decreasing concentration: an ionizable lipid, with an apparent pKa typically between 6 and 7, which encapsulates the RNA payload and effects intracellular delivery; cholesterol, for modification of zwitterionic lipid and ionizable lipid fluidity (melting temperatures); a zwitterionic helper lipid, which forms low-permeability surface bilayers on processed LNPs; and a PEG-lipid, the steric stabilizer to enhance the LNP stability at neutral pH^7^. The formulation of LNPs relies on antisolvent nanoprecipitation through rapid and controlled mixing of the ethanolic lipid solution (solvent) and the acidic buffered RNA solution (antisolvent). Homogenous fluid mixing triggers lipid precipitation, as the four lipid components (three amphiphiles and one hydrophobic cholesterol) are supersaturated species, which assemble into LNPs following a nucleation and particle growth mechanism. The hydrophilic RNA cargo forms an electrostatically complexed hydrophobic salt with the ionizable lipid component and enters the growing lipid-RNA aggregates. LNP assembly is usually performed at an acidic pH (pH 3 to 4), which charges the ionizable lipid and transforms it from a hydrophobic oil into an efficient cationic surfactant that can readily form the RNA-lipid complex required for RNA encapsulation. The large, cationic surface charge of lipid assemblies at the low assembly pH generates a long-ranged electrostatic repulsive force while the PEG-lipid surface density remains low. Exchanging this acidic-buffered, mixed solvent for a neutral aqueous buffer raises the pH above the ionizable lipid pKa, lowering the surface charge such that electrostatic repulsion is insufficient to prevent LNP aggregation. However, LNPs do not aggregate into macroscopically phase-separated materials because of the presence of the PEG-lipid. Every aggregation event reduces the specific surface area of the particle and “concentrates” the PEG-lipid on the surface until the PEG achieves a sufficient density to resist further particle-particle aggregation. LNPs made with intermediate pH buffers (pH 5-6) or very high ionic strengths display hybrid behaviors, with both the electrostatic and steric stabilization mechanisms (“electrosteric” describing the sum of these two stabilizing forces). Repulsive electrosteric forces at the LNP surface effectively arrest further particle growth and mark completion of the initial self-assembly process at some critical, characteristic size depending on the composition of both the organic lipid and aqueous RNA solutions, including pH and salts^8,9^. Given the dependence of this process on supersaturation and interfacial fluid gradients, LNP assembly is inherently influenced by mixing conditions including homogeneity of fluid mixing and local solvent compositions during nanoprecipitation^10^. The presence of the several amphiphilic species, and the ion pairing between the ionizable lipid and RNA, further complicate the self-assembly process. A recent study modeling LNP self-assembly via a population balance approach relied on a simplifying assumption that all lipid species in the system are purely hydrophobic and have the same solubility as cholesterol, a 38.5 mol% component; this does not capture the amphiphilicity of 61.5 mol% of the lipids in this system, which is difficult to model^5^. We give this as an example to motivate the need for experimental measurements on this complex process.

## Experimental Section

### Materials

ALC-0315 (6-((2-hexyldecanoyl)oxy)-N-(6-((2-hexyldecanoyl)oxy)hexyl)-N-(4-hydroxybutyl)hexan-1-aminium), DSPC (1,2-distearoyl-sn-glycero-3-phosphocholine), cholesterol, and DMG-PEG2000 (1,2-dimyristoyl-rac-glycero-3-methoxypolyethylene glycol-2000) were purchased from Avanti Polar Lipids (Alabama, USA). Purified Torulla Ambion yeast RNA (yRNA) was purchased from ThermoFisher Scientific (Massachusetts, USA). Polyadenylic acid potassium salt (polyA) was purchased from Sigma Aldrich (Missouri, USA). Absolute ethanol (99.8%), Tetrahydrofuran (THF, 99.8% unstabilized) and buffer salts (sodium acetate) were purchased from Fisher Scientific (Massachusetts, USA) and FisherSci (Sweden). Polycaprolactone-b-poly(ethylene glycol) (PCL5k-b-PEG5k) and Poly(e-caprolactone) homopolymer (PCL – avg. Mw: 12 kDa) were purchased from PolymerSource (Montreal, Canada). Sodium polystyrene sulfonate standard calibration kit was purchased from Scientific Polymer Products Inc. (New York, USA). Ultrapure (UP) water was procured using an in-house water filtration system (ThermoFisher Scientific AquaPro, dispensed at 18.2 MΩ -cm resistivity).

## Methods

### Lipid and polymer nanoparticle formulation and process parameters

RNA-LNPs and empty LNPs (eLNPs) were produced by Flash NanoPrecipitation achieved through mixing of an ethanolic stream containing dissolved lipids and an aqueous stream containing 20 mM sodium acetate buffer solution (pH 5.0) with one of polyA, yeast RNA or no RNA, respectively (0.6 mg/mL RNA when present). The lipid stream contained a 12 mg/mL mixture of ALC-0315 (7.07 mg/mL), DSPC (1.46 mg/mL), cholesterol (2.75 mg/mL), and DMG-PEG2000 (0.68 mg/mL) at a molar ratio of 50/10/38.5/1.5 mol%, respectively. The loaded LNP formulations had an N/P = 5, where N/P is defined as the molar relationship between the ionizable lipid nitrogen (N) and RNA phosphate group (P). eLNPs were produced in the same manner using the same sodium acetate buffer solution (20 mM, pH 5.0) without RNA. Mixing was carried out in either a CIJM, MIVM^4,11^, or T-mixer with aqueous and organic feed streams delivered using syringe pumps (PHD Ultra - Harvard Apparatus, MA, USA) under variable operational total flow rates (TFRs) and aqueous:organic flow rate ratios (FRRs) presented in **Table 1**.

**Table 1.**
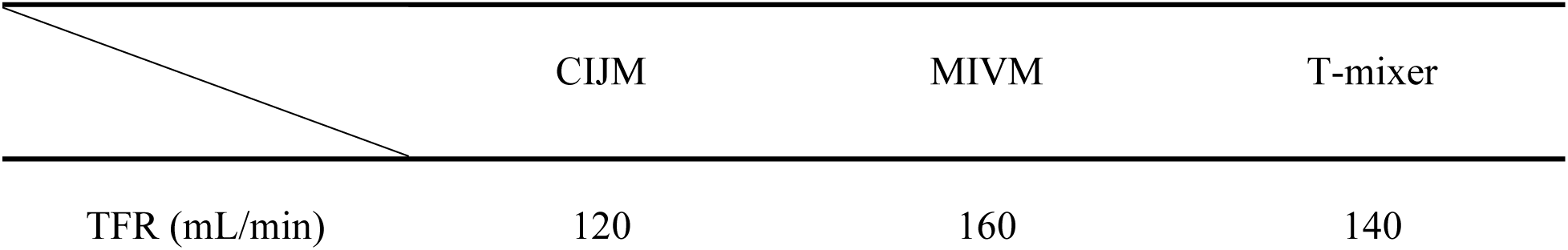
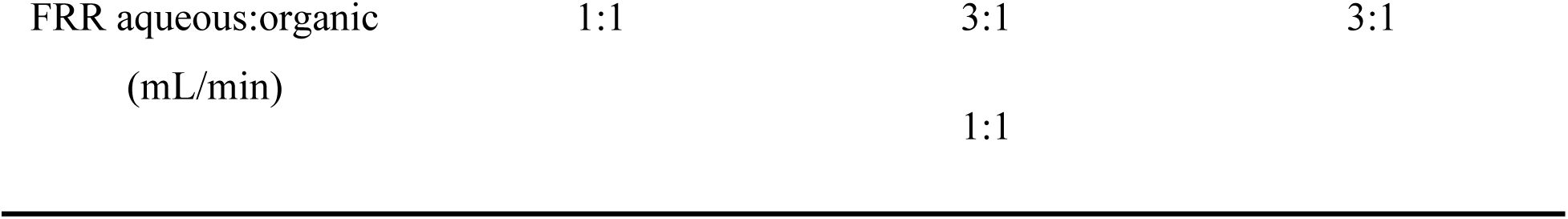
Operational TFRs and FRRs for the mixers employed in the kinetics measurements.

The two PNP formulations studied were also formed by Flash NanoPrecipitation using a CIJM or an MIVM. Small PNPs of ∼30 nm.r (intensity-weighted hydrodynamic radius measured by DLS) were produced in a 4-inlet MIVM, where the organic feed stream consisted of 5 mg/mL PCL homopolymer and 5 mg/mL PCL-b-PEG stabilizer dissolved in THF. The organic feed stream was mixed with UP water in 3:1 aqueous:organic FRR using an MIVM at a TFR of 160 mL/min. Large PNPs of ∼75 nm.r were produced in a two jet CIJM, where the organic feed stream consisted of 15 mg/mL PCL and 5 mg/mL PCL-b-PEG dissolved in THF. The organic feed stream was mixed with UP water in a 1:1 FRR using CIJM at a TFR of 120 mL/min. For lab-based size validation, the effluent streams of each mixer were quenched in UP water, by directing the mixer effluent stream into a vial containing the necessary amount of UP water to reach a final 10% v/v THF concentration, to kinetically arrest solubility driven PNP size evolution.

### Physicochemical characterization of LNPs and PNPs

Particle size distributions (hydrodynamic) and PDI were determined using dynamic light scattering (DLS) on a Malvern Zetasizer Pro (Malvern Panalytical, United Kingdom), at a 173° scattering angle. Temperature and acquisition settings were set to 25°C and auto, respectively. Measurements were performed in triplicate, and all samples were diluted 10-fold in buffer (LNPs) or UP water (PNPs) prior to analysis. All LNP samples produced at an FRR of 3:1 Aq:Org were measured without prior dilution, with the measurement parameters adjusted to account for the increased viscosity and refractive index associated with 25% v/v of ethanol. LNP samples produced at a 1:1 FRR were initially quenched to 10% v/v of ethanol and subsequently diluted to 1% v/v of ethanol prior to DLS analysis. Both PNP samples produced at an FRR of 1:1 and 3:1 aqueous:organic were diluted to a final THF fraction of 1% v/v prior to DLS analysis, owing to the incompatibility of THF with plastic cuvettes.

### Flowthrough synchrotron small-angle X-ray scattering

For experiments at MAX IV:

Flowthrough SAXS experiments were carried out in air in transmission mode at the CoSAXS beamline at the 3 GeV ring at MAX IV laboratory (Lund, Sweden). Samples were passed through a thin-walled quartz capillary with a 1.5 mm inner diameter and a wall thickness of 10 microns (Hilgenberg GmbH, Malsfeld, Germany). Frames were acquired using a fixed X-ray energy of 12.4 KeV (λ = 1.00 Å) with a hybrid pixel X-ray detector Eiger2 4 M (DECTRIS). The sample to detector distance (SDD) was set to 3.5 m with silver behenate (AgBeh) as calibration standard to determine the *q-*scale (∼0.003 – 0.309 Å^-1^) where *q* denotes the scattering vector, defined as:

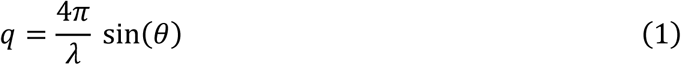

Where λ is the X-ray wavelength and θ is the scattering angle. The *q* positions of scattering peaks were used to determine the d-spacings of liquid crystalline domains (internal ordering) in the lipid nanoparticles, using the following relationship:

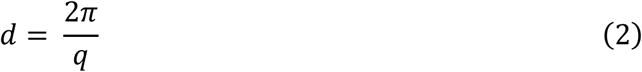

2D detector images were radially and azimuthally integrated to obtain 1D scattering curves using the MatFRAIA algorithm as implemented in Python^12^.

For experiments at the Advanced Photon Source (APS):

SAXS experiments were carried out in air in transmission mode at the 12-ID-B beamline of the Advanced Photon Source, Argonne National Laboratory (IL, USA). Samples were flowed through a thin-walled quartz capillary with a 1.5 mm inner diameter and a wall thickness of 10 microns (Charles Supper, MA, USA) in an in-house fabricated flow-cell. Frames were acquired using a fixed X-ray energy of 13.3 KeV (λ = 0.932 Å) and an Eiger2 9 M X-ray detector (DECTRIS). The sample to detector distance (SDD) was set to 3.6 m with AgBeh as calibration standard to determine the *q-*scale (∼0.002 – 0.48 Å^-1^).

In all flowthrough experiments, X-ray exposure time was set to 50 ms per frame with a total of 300 frames per acquisition (15 seconds of fluid flow). The initial 50-80 frames (3-4 s) and the final 20 frames (1 s) were omitted from preprocessing and subsequent data analysis, as these intervals were designated as pre- and post-waste (i.e., periods of start-up or shut-down flow associated with poor mixing, in which particle characteristics may differ). Background scattering patterns with the relevant solvent composition were collected before the start of each sample run, under identical exposure conditions. Between runs, the capillary was flushed with water and ethanol or THF (LNP or PNP samples, respectively) for cleaning. All LNP samples had a total lipid and RNA concentration of 3 mg/mL and 0.15 mg/mL, respectively, at the point of X-ray exposure. PNP samples with an average size (R_h_) of ∼30 nm.r and ∼75 nm.r had a total solids concentration of 2.5 mg/mL and 7.5 mg/mL, respectively, at the point of X-ray exposure. Both lipid and polymer organic feed streams were prepared on-site at each beamline immediately prior to the flowthrough SAXS experiment. For static measurements of fully equilibrated LNPs, 20 frames were acquired at an X-ray exposure of 50 ms.

### SAXS data analysis

SAXS frames were averaged and buffer subtracted using the opensource software BioXTAS RAW^13^. Subsequent data fitting of a pair-distance distribution function *p*(r) was carried out using GNOM^14^. The *p*(r) distribution provides information on changes in real-space dimensions of the particles through indirect Fourier transformation of the 1D SAXS profile. For PNPs at a 75 nm.r R_h_, the fit was applied to a *q*-range of 0.0022 – 0.30 Å^-1^, which includes all measured low-*q* data points. Sensitivity analysis (**Supporting information**) showed that setting *q*-max ranging from 0.30 – 0.488 Å^-1^ for these profiles did not significantly alter extracted R_g_. For all LNPs (FRR 3:1 and 1:1) as well as 30 nm.r R_h_ PNPs, the fit was applied to a *q*-range of 0.003 – 0.31 Å^-1^.. Sensitivity analysis (**Supporting information**) showed that setting *q*-min ranging from 0.002 – 0.0048 Å^-1^ affected the extracted R_g_ (up to 14%). To minimize the influence of the initial points low-*q* data on the *p*(r) analysis, the initial data points were excluded from the data collected at APS, while the full q-range was used in the MAX IV data. Accordingly, a consistent fitting range was applied to facilitate qualitative comparison between runs within the same condition. The fits were adjusted manually to optimize quality of the fit (minimizing χ^2^), total estimate and residuals distribution, while ensuring that the distribution profile exhibited a physically meaningful decay towards zero at D_max_.

As the SAXS measurements probed particles directly during in-line nanoprecipitation at flow rates of up to 160 mL/min, the scattering volume inevitably contained heterogeneous particle populations, including transient larger particle species. Consequently, *p*(r)-derived *D*_max_ values were particularly sensitive to these species and the average sizes extracted exceeded the dimensions of the dominant colloidal population. *D*_max_ was therefore not interpreted as a quantitative particle diameter. Instead, changes in the *p*(r) distributions were used qualitatively to follow changes in averaged characteristic particle dimensions during particle formation. Due to the heterogenous nature of LNP shape and electron density distribution, the obtained R_g_ are not directly comparable to the R_h_ obtained using DLS. The R_g_ obtained from SAXS derived *p*(r) was compared to the hydrodynamic radius distributions obtained from DLS analysis only as a qualitative assessment of particle dimensions.

### Experimental configuration for flowthrough SAXS at the CoSAXS and 12-ID-B beamlines

To resolve millisecond-scale kinetics of nanoparticle formation, the TFR of the effluent stream of the mixer must be sufficiently high and the mixer plus time-of-flight controlling tubing volume (V) sufficiently small for material to advect into the SAXS window at the target time (t = V/ TFR). Therefore, the configuration used in the current study relied on high-throughput turbulent mixers capable of delivering flows of up to 160 mL/min, while still maintaining favorable LNP physicochemical properties such as size, PDI and EE%.

Using turbulent flow mixers in the experimental configuration served several purposes in the current study. Mainly, it was the aim of this study to observe LNP formation kinetics that were driven primarily by diffusion limited aggregation (DLA) of supersaturated lipids, rather than limited by solvent mass transport in the case of slower mixing. The turbulent flow mixers employed in this study are designed to satisfy the condition of t_mix_ < t_assembly_ which implies a low Damköhler number, Da = t_mix_/t_assembly_ < 1, whereby mixing occurs on much shorter timescales than particle self-assembly^15^. Additionally, high-throughput turbulent mixers are industrially relevant models, which have been demonstrated to be feasible at scale for mRNA-LNP vaccine production16, where information on formation kinetics can prove valuable for manufacturing process design.

**Figure 1** shows the experimental configuration of the three turbulent flow mixers applied in this study: an MIVM (Holland Applied Technologies, IL, USA) setup at the CoSAXS beamline (MaxIV, Lund, Sweden), a CIJM (Holland Applied Technologies, IL, USA) and an in-house fabricated T-mixer (1 mm ID) setup at the 12-ID-B beamline (APS, IL, USA). **Figure 1e** shows a bespoke CIJM with a quartz window (1 mm) mounted on a 3D-printed resin holder, allowing acquisitions at the point of mixing (12 ms - TFR 120 mL/min). For acquisitions at longer residence times, tubing (3.175 mm OD x 1.58 mm ID - PEEK - Cole-Parmer, IL, USA) of varying lengths were connected between the outlet of the mixer (MIVM and CIJM) and the inlet of the quartz capillary flow-cell (1.5 mm ID). This allows for SAXS acquisitions at what would be milliseconds after nanoprecipitation, building on our previously published setup^4^. Importantly, in this work, the fluid residence times (LNP age from onset of mixing) should not be confused with X-ray exposure times defined in the SAXS methods (50 ms). In the flowthrough SAXS experiments, the fluid in the capillary was under continuous steady flow, where each experimental acquisition generates frames from the same exposure point on the capillary, in which successive fluid elements pass through the X-ray beam. It is the geometry of the mixer, the delay tubing at the outlet and the volumetric flowrate that define the fluid residence time at the exposure point.

**Figure 1.**
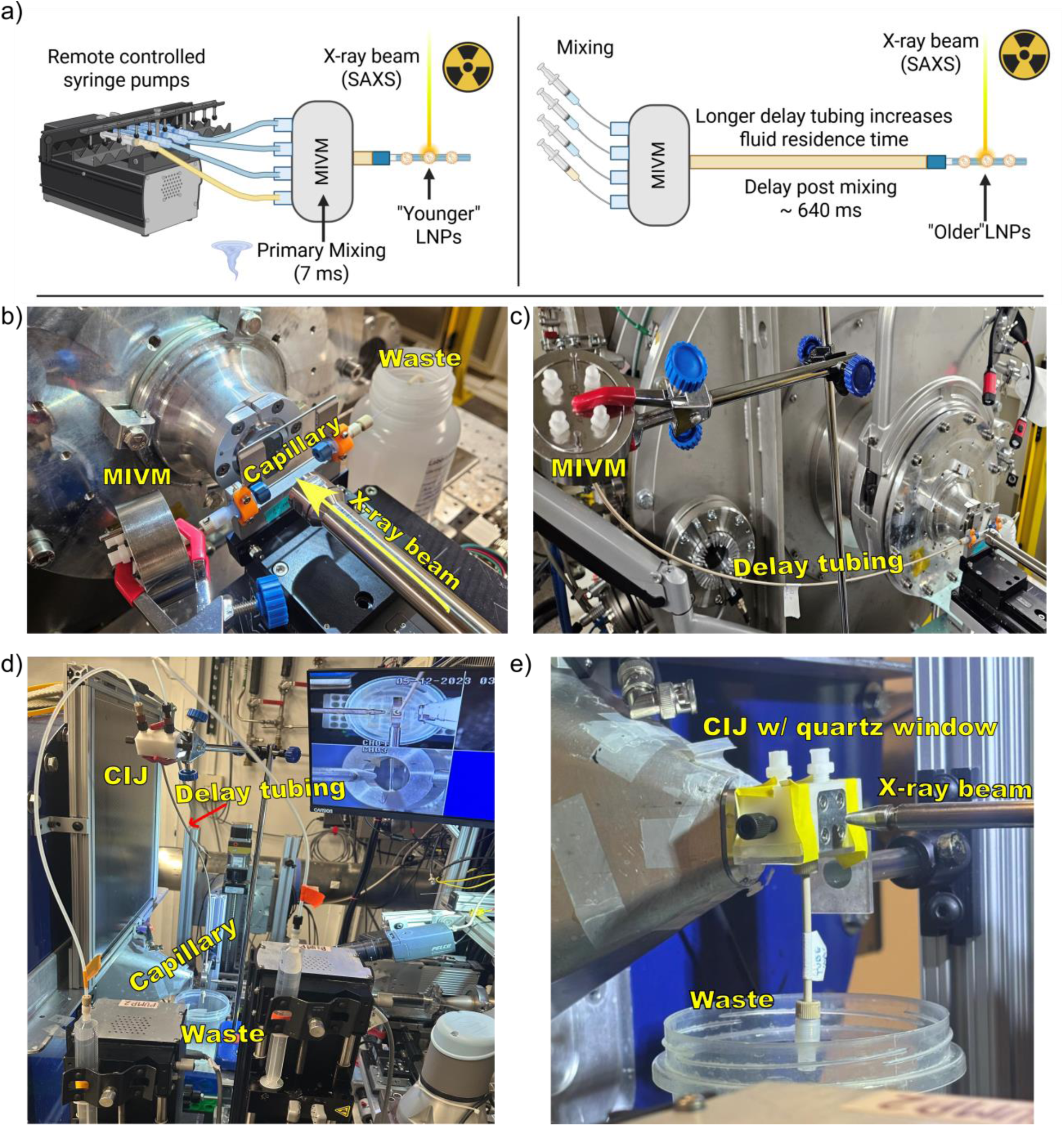
a) Schematic representation of experimental configuration in the beamline sample environment, including description of delay tubing that allows acquisition at defined residence times. b) Labelled image of MIVM-SAXS experimental configuration at the CoSAXS beamline, showing the MIVM immediately in-line with the quartz capillary for data acquisition at ∼45 ms residence time. Buffer and ethanol streams were delivered to the MIVM using a remotely controlled syringe pump setup at a TFR of 160 mL/min and an Aq:Org FRR of 3:1. c) Labelled image showcasing an example of the MIVM-SAXS configuration with the longest delay tubing for data acquisition at ∼640 ms residence time at a TFR of 160 mL/min and an Aq:Org FRR of 3:1. d) Labelled image showcasing the CIJM-SAXS setup with the longest delay tubing for data acquisition at ∼405 ms residence time at a TFR of 120 mL/min and an Aq:Org FRR of 1:1. e) CIJM mixer with a quartz window (1 mm) aligned with the X-ray beam for frame acquisition at the point of mixing (12 ms at 120 mL/min TFR).

### Sample preparation for single particle Raman analysis

Single particle automated Raman trapping analysis (SPARTA) measurements were performed using an AGIS I (SPARTA Biodiscovery; London, UK) spectrometer. Backgrounds were collected on a 10 mM, pH 7.4 (range 7.3 - 7.4) HEPES buffer with a 15 second exposure time. Particle samples in the same HEPES buffer were analyzed by diluting to 0.2 - 0.6 mg/mL total solids (approximately 0.25 - 1.0 x10^12^ particles/mL for a ∼100 nm diameter LNP) with a 15 second exposure time and 1 second trap reset time. LNPs at this concentration and an adjusted trapping threshold (typically in the range of 11 – 13) yielded an average trapping iteration number in the range of 2 – 5, indicating successful capture of single, random particles each time the particle trap re-engaged after the 1 second delay window. Raman spectra were processed using SPARTA Discovery software to correct instrument background, exclude outlier spectra, and create the final averaged, normalized spectra presented here. The Raman spectrum of an LNP is dominated by the lipid components, as expected given the majority of the LNP mass is lipids, but the PEG stabilizer and RNA cargo are difficult to quantify in the Raman spectrum. The low Raman intensity of the RNA cargo, due to low concentration and an intrinsic small Raman scattering coefficient, presents a challenge in directly measuring the untagged cargo. We instead prepared LNPs with poly(styrene sulfonate) (PSS) as a linear polyanion mimic of the RNA cargoes used in the time resolved SAXS measurements. PSS-containing LNPs were prepared using a CIJM in the same fashion as described above, with two minor changes. The lipid feed stream concentration was 6 mg/mL in ethanol at the same molar ratio of ALC-0315 (3.53 mg/mL), cholesterol (2.75 mg/mL), DSPC (1.46 mg/mL), and DMG-PEG2000 (0.36 mg/mL). This lipid stream was mixed against an aqueous stream with a 20 mM, pH 4 sodium acetate buffer containing either 318 µg/mL for *in situ* loaded PSS-LNPs or 0 µg/mL for eLNPs. eLNPs were *post hoc* loaded by mixing the eLNPs against a 318 µg/mL solution of PSS in the same acetate buffer. Note that at this PSS concentration, the N:P ratio is 3 rather than the 6 used with RNA samples above. This increase in PSS loading was chosen to maximize PSS signal in the SPARTA measurements without excessive increase in the particle hydrodynamic diameters. All PSS-LNPs were dialyzed against the 10 mM, pH 7.4 HEPES buffer used for SPARTA measurements. Dialysis tubing was 6 – 8 kDa MWCO RC1 regenerated cellulose tubing (Repligen; Waltham, MA) with a retentate volume of 5 mL in an external media volume of 500 mL. Each dialyzed sample was equilibrated for 4 hours, with one change of the external media at the 2-hour mark. DLS measurements were performed on dialyzed PSS-LNP samples without additional dilution. *In situ* loaded polyA LNPs were also synthesized and dialyzed for SPARTA measurement as a control for LNPs without PSS cargo. A reference spectrum for solid PSS (504,400 Da) was collected with an air background and an exposure time of 3 seconds with the trapping threshold set to its minimum value to capture all spectra.

## Results and discussion

### Time-resolved flowthrough SAXS captures millisecond scale particle formation kinetics in turbulent mixing

To validate the suitability of this setup for tracking nanoparticle formation and structural evolution, an initial experiment was conducted using polymeric nanoparticles (PNPs). The PNP formulations chosen for this step consist of a hydrophobic core component (PCL) and an amphiphilic stabilizer component (PCL-b-PEG). These PNP systems form a core-shell structure with a hydrophobic homopolymer (PCL) core and a surface-adsorbed layer of amphiphilic block copolymer (PCL-b-PEG) which imparts the stabilizing PEG polymer corona. PCL polymer coils rapidly collapse^17^ in the mixed solvent-antisolvent system, yielding collapsed PCL globules. Homopolymer PCL globules aggregate upon collision, leading to a DLA assembly of the core. Block copolymer PCL-b-PEG has a collapsed PCL globule attached to a soluble PEG coil, which adsorbs to the growing PCL core in a reaction limited aggregation (RLA) mechanism because only PCL-PCL collisions result in block copolymer adsorption; collisions between the PEG coil and the PCL core yield no stabilizing block copolymer adsorption. The ratio between the PCL and PCL-b-PEG can be tuned to yield differently sized PNPs, with different overall assembly kinetics reflecting the rates of PCL homopolymer core growth by DLA and PCL-b-PEG stabilization by RLA^18^. A PNP formulation yielding a particle size distribution (PSD) centered around a 30 nm.r intensity-weighted hydrodynamic radius (nm.r) was selected as a proxy for primary LNPs, which typically display a particle radius in the 15 - 30 nm range^8^. These small particles should assemble within milliseconds of mixing and would therefore be fully formed at all acquisition times accessible in this experimental setup. The 30 nm.r PNPs were analyzed using the MIVM-SAXS setup imaged in **Figure 1b** and **1c**, at a TFR of 160 mL/min and an FRR of 3:1. Larger PNPs with an average size of 75 nm.r were analyzed using the CIJM-SAXS setup shown in **Figure 1d** and **1e**, at a TFR of 120 mL/min and an FRR of 1:1. Compared with the 30 nm.r particles, these particles were expected to assemble on longer timescales^18^, such that their evolution could be observed within the temporal resolution of the SAXS setup. **Figure 2** shows the results of the setup suitability test. In **Figure 2a** and **Figure 2c**, the averaged SAXS frames are plotted for each acquisition timepoint for 75 nm.r PNPs and 30 nm.r PNPs, respectively. The frames show clear low-*q* particle scattering at all measured timepoints, with amorphous particle cores evidenced by a featureless high-*q* region. The *p*(r) function was obtained through indirect Fourier transformation of the scattering profiles to provide a shape-independent real-space representation of the particle morphology. The resulting *p*(r) plots are presented in **Figure 2b** and **2c**. In the case of the 30 nm.r PNPs particles, the averaged frames from SAXS acquisitions at each time point result in similar *p*(r) profiles that exhibit a near-spherical morphology. In comparison, the profiles extracted from the averaged frames of the 75 nm.r PNPs initially exhibit a similar p(r) profile, also suggesting a spherical morphology at 12 ms and consequently exhibit a more skewed p(r) profile, suggesting a more polydisperse mass distribution at the later probed timepoints. This is potentially explained by the maturation of the PEG shell that forms around the particle core, as well as structural relaxation after nanoprecipitation, which may contribute to a change in the electron density distribution and deviation from a monodisperse spherical *p*(r) profile.

**Figure 2.**
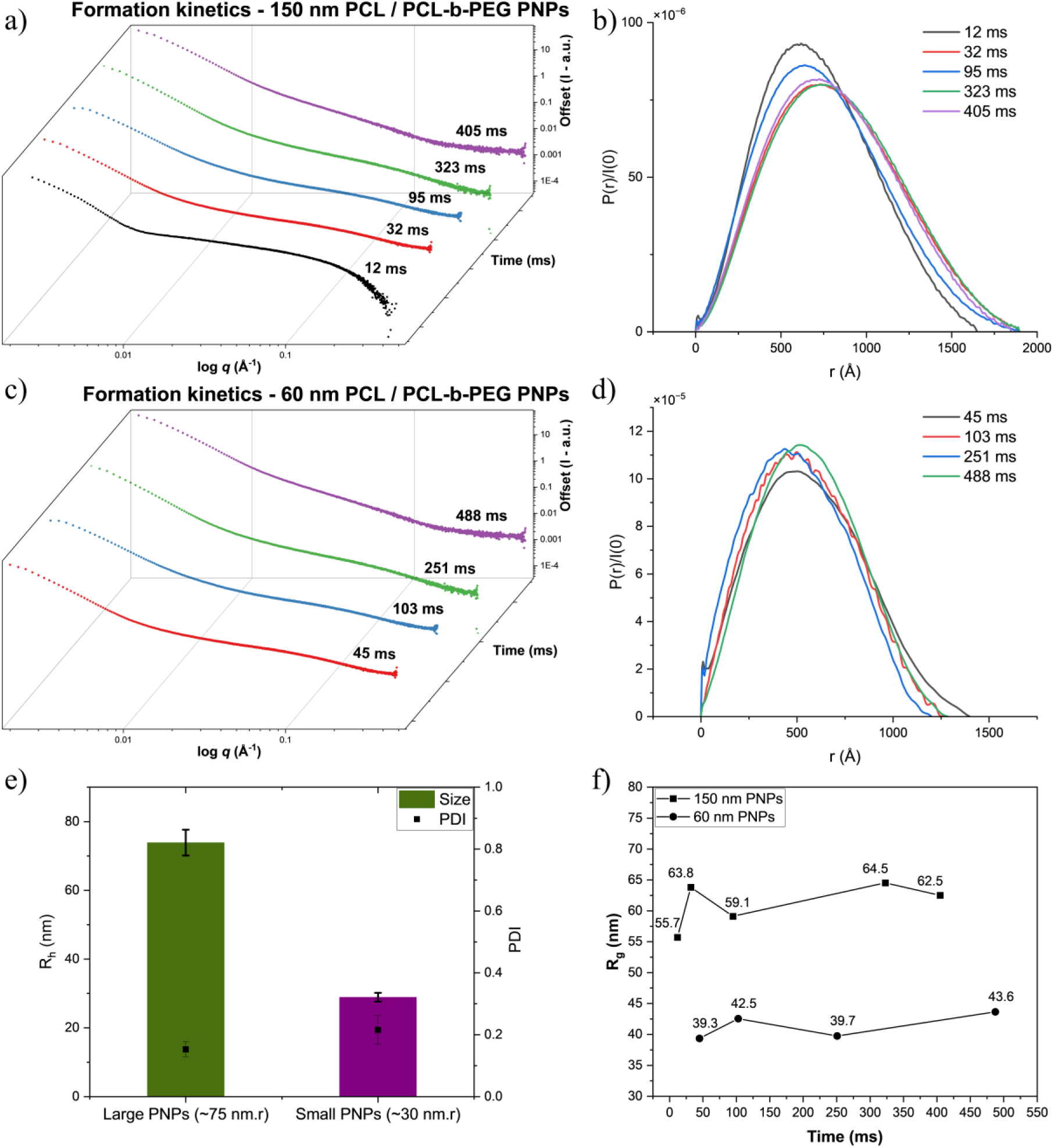
a) Flowthrough CIJM-SAXS measurements for PCL/PCL-b-PEG PNPs displaying an average R_h_ of 75 nm. Measurements were carried out at a fluid residence time of 12, 32, 95, 323, and 405 ms post-mixing at a TFR of 120 mL/min and an FRR of 1:1 Aq:Org. b) *p*(r) profiles for each averaged scattering pattern of the flowthrough CIJM-SAXS measurements of 75 nm.r PNPs in Fig 2a. c) Flowthrough MIVM-SAXS measurements for PCL/PCL-b-PEG PNPs displaying an average R_h_ of 30 nm. Measurements were carried out at a fluid residence time of 45,103, 251, and 488 ms post-mixing at a TFR of 160 mL/min and an FRR of 3:1 Aq:Org. c) *p*(r) profiles for each averaged scattering pattern of the flowthrough MIVM-SAXS measurements of 30 nm.r PNPs in Fig 2c. e) DLS measurements showing the intensity weighted z-average hydrodynamic radius R_h_ and PDI values for the fully formed 75 nm.r and 30 nm.r PNPs. (n = 3, ±SD). f) R_g_ (nm) extracted from each timepoint for both 75 nm.r and 30 nm.r PNPs plotted against fluid residence time (ms).

A radius of gyration (R_g_) was extracted from the *p*(r) of each scattering curve for both 75 nm.r and 30 nm.r PNPs and plotted against time in **Figure 2f**. The R_g_ values extracted for 30 nm.r particles show similar dimensions at all probed timepoints, suggesting that no overall evolution in particle size occurs after the fluids are mixed in and exit the MIVM. The 30 nm.r particles can be considered fully formed immediately downstream of the MIVM by 45 ms (the earliest probed timepoint), exhibiting a size that is comparable to the corresponding hydrodynamic radius (R_h_) obtained using DLS, for the fully formed PSD, presented in **Figure 2e**. For the 75 nm.r PNPs, the apparent *R_g_*was lower at the earliest measured time point (12 ms) than at later time points, although the values subsequently fluctuated around an R_g_ of 59-65 nm. This may indicate changes in the characteristic dimensions of the scattering population following mixing; however, the magnitude of these changes should be interpreted cautiously given the influence of particle heterogeneity and larger species on the *p*(r) analysis. The CIJM used for the 75 nm.r particles enables observation of this earlier timepoint than was possible in the MIVM, and the larger particles are ∼16x more massive than the smaller particles are which requires a correspondingly longer assembly time which can be detected with our time-resolved SAXS setup. Interestingly, these differences were also visible in the SAXS capillary, whereby the earliest timepoints were less turbid at the point of X-ray exposure, which implies the evolution of larger sizes between 12 to 405 ms is a real phenomenon which increased the *visual* light scattering as well. Despite the limitations associated with quantitative size determination from the *p*(*r*)analysis, the measurements demonstrate that the flow-through SAXS setup can reproducibly probe particle populations at defined time points following mixing and clearly distinguish between particles of different characteristic dimensions (60 nm vs 150 nm PNPs). Overall, the results suggest that the mixer-SAXS platforms can capture relative millisecond scale evolution of average nanoparticle size over time (previously this setup had only been used to probe the time scales of liquid crystalline mesophase formation, at higher *q*)^4^.

Given that the 30 nm.r PNP core formed rapidly through DLA, it was subsequently investigated whether LNPs would exhibit similar formation behavior. In contrast to the aggregation of PNPs consisting of PCL, which is a purely hydrophobic, polymeric core material, three out of four lipid small molecule components of an LNP are amphiphilic. The question remained whether the amphiphilicity imparted an orientational dependency for the successful collisions of the lipid molecules during self-assembly, and whether this would contribute to a delay of particle maturation by shifting particle core assembly from a rapid DLA to a slowed RLA fashion.

### LNP formation is completed within 17 ms across multiple nucleic acid cargo lengths at a 3:1 flow rate ratio

To probe the formation kinetics of LNPs, the applied MIVM-SAXS setup was extended to investigate LNPs encapsulating yeast RNA (yRNA) and polyA, as well as eLNPs, all produced at 3:1 FRR at a TFR of 160 mL/min for continuous nanoprecipitation. **Figure 3** shows the results of the MIVM-SAXS analysis at various timepoints after mixing. In all measured *in situ* nanoprecipitations, the RNA-loaded LNPs were produced in-line with the quartz capillary by mixing an aqueous stream of 0.6 mg/mL yRNA or polyA in a 13.32 mM acetate buffer (pH 5.0) against a 12 mg/mL ethanolic lipid stream at a 3:1 FRR and a TFR of 160 mL/min or 140 mL/min using either the MIVM or T-mixer, respectively. eLNPs were produced in a similar manner using buffer without dissolved RNA. **Figure 3a** shows the averaged SAXS profiles for each timepoint measured for yRNA-loaded LNPs. At the earliest measured timepoint, 45 ms, the averaged profile shows clear low-*q* particle scattering as well as the appearance of a correlation peak at ∼ 0.15 Å^-1^ in the high-*q* region, indicating the formation of an internal liquid crystalline (LC) phase within the particle core, consisting of ionizable lipid-RNA complexes at the interface of lipid rich regions and water channels^19^. The shape of the scattering profile did not change significantly when measured at longer fluid residence times. The same trend was also observed in the scattering profiles of polyA-loaded LNPs (**Figure 3c**). While both model RNAs include a heterogenous blend of nucleic acid strand lengths, polyA molecules are on average significantly (∼10x) larger than yRNA strands that had an approximated average MW of 6500 g/mol. Due to the size differences of the cargo, it may be expected that encapsulation of the larger polyA RNA would result in longer self-assembly times or delayed appearance of an LC peak. This was not observed across the timescales studied here. The *p*(r) profiles of the flowthrough scattering patterns recorded in **Figure 3a** and **3c** showed an elongated morphology evident by the tailing of the bell-shaped curve towards D_max_ (**Figure 3b** and **3e**). The prolate spheroid (aspect ratio typically ∼ 1.0 - 1.2) morphology, extracted from the *p*(r) profile, has been previously reported for LNPs^20^. This contrasts with the breadth of existing cryo-TEM micrographs across multiple studies reporting that LNPs largely exhibit a roughly spherical morphology, albeit without direct measurement of the particle dimensions^21^. It is therefore important to mention that given the reported heterogeneity in size, shape and cargo loading within LNP dispersions^22–24^, it is likely that the *p*(r) profile of the analyzed particle systems reflects an averaged shape arising from this heterogeneity (and polydispersity), rather than being reflective of a single, universal particle shape. SAXS is not a single particle analysis technique: the large (compared to a single nanoparticle) irradiated volume by the focused beam and time-integrated signal of each frame make it an obligate ensemble measurement. This is particularly relevant for bulk measurements of non-fractionated LNPs formed *in situ*, where scattering signal intensity fluctuates over time as heterogenous particle populations and occasional larger particles pass through the beam cross-section. The patterns plotted in **Figure 3a** and **3c** represent ensemble-averaged frames collected over several seconds of running the mixer at a flowrate of 160 mL/min. Consequently, the measured profiles reflect the collective scattering contribution of a heterogenous particle population, with occasional aggregates potentially biasing the averaged scattering patterns toward larger length scales. This variation in particle R_g_ within an LNP formulation was recently highlighted in a study characterizing colloidal subpopulations of LNPs separated by asymmetric flow field flow fractionation (AF4) and coupled to SAXS^25^. The R_g_ values extracted from the *p*(r) fits for both yRNA and polyA loaded LNPs lies within the PSD obtained from the DLS measurement of the fully formed particles in 25% EtOH (**Figure 3h**). This average particle size does not evolve over the course of the fluid residence times probed (up to 488 ms). Additionally, the high-*q* correlation peak present at the earliest timepoints suggest that nanostructured “core” (cryo-TEM images for LNPs at pH 5.0 in **Figure S2**) lipid aggregates are formed by the time the LNPs exit the mixer. As the core occupies the interior of the particle, it likely functions as a nucleus or template for the development of the surrounding shell. Therefore, in the context of LNPs and the data presented in **Figure 3**, the formation of this internal core will precede DSPC bilayer enveloping and DMG-PEG2000 PEG-lipid surface deposition ^26^, in a well-mixed system.

**Figure 3.**
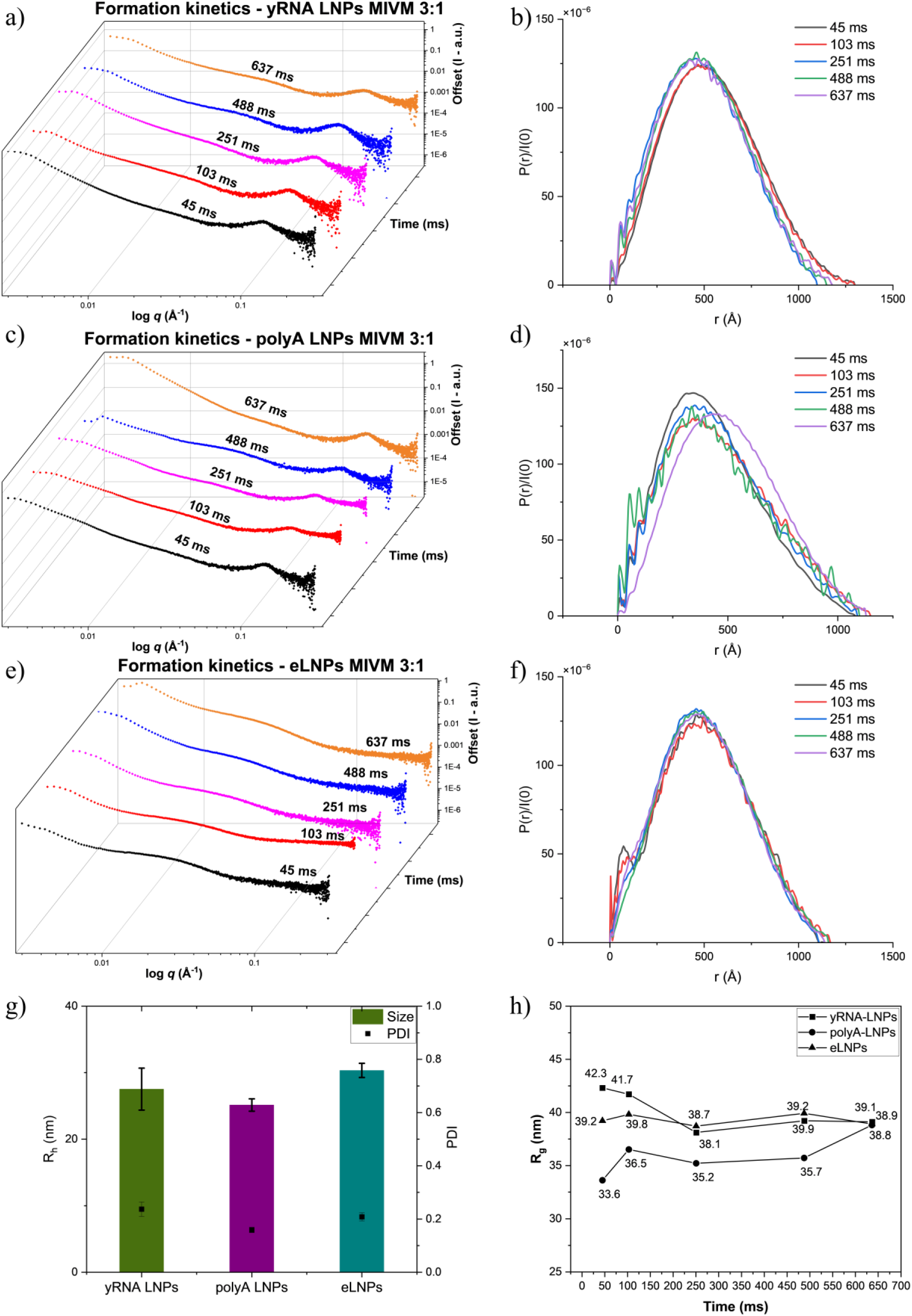
a) Flowthrough MIVM-SAXS measurements for *in situ* nanoprecipitation of yRNA-loaded LNPs. Measurements were carried out at a fluid residence time of 45, 103, 251, and 488 ms post-mixing at a TFR of 160 mL/min and an FRR of 3:1 Aq:Org. b) *p*(r) profiles for each averaged scattering pattern of the flowthrough MIVM-SAXS measurements of yRNA-loaded LNPs in Fig 3a. c) Flowthrough MIVM-SAXS measurements for *in situ* nanoprecipitation of polyA-loaded LNPs. Measurements were carried out at a fluid residence time of 45, 103, 251, and 488 ms post-mixing at a TFR of 160 mL/min and an FRR of 3:1 Aq:Org. d) *p*(r) profiles for each averaged scattering pattern of the flowthrough MIVM-SAXS measurements of polyA-loaded LNPs in Fig 3c. e) Flowthrough MIVM-SAXS measurements for *in situ* nanoprecipitation of eLNPs. Measurements were carried out at a fluid residence time of 45, 103, 251, and 488 ms post-mixing at a TFR of 160 mL/min and an FRR of 3:1 Aq:Org. f) *p*(r) profiles for each averaged scattering pattern of the flowthrough MIVM-SAXS measurements of eLNPs in Fig 3e. g) DLS measurements showing the intensity weighted z-average hydrodynamic radius R_h_ for the fully formed loaded and eLNPs (n = 3, ±SD). h) R_g_ (nm) extracted from each timepoint for yRNA loaded, polyA loaded and eLNPs plotted against fluid residence time (ms).

Therefore, the results suggest that both yRNA and polyA LNPs are fully formed by 45 ms and do not undergo further self-assembly beyond this point, excluding structural rearrangements associated with downstream processing steps such as buffer exchange and pH shifts. This conclusion is dependent on the criteria chosen to define “fully formed” LNPs. Here, we adopted a conservative definition that required an average particle size, extracted from the *p*(r), which does not evolve over the probed timeframe, and which lies within the PSD measured by DLS (**Figure 3g**), as well as the presence of an LC phase peak with a position and shape consistent with that observable in fully equilibrated LNPs measured *ex situ*, which does not evolve over the measured interval. For each residence time condition, consecutive SAXS frames were acquired during a continuous flowthrough experiment and the frames averaged for improved statistics. The averaged frames were then subject to data analysis as outlined in the SAXS data analysis section. Independent experimental replicates were not measured for each specific condition, owing to the substantial resource consumption associated with running lipids and RNA through a high-volume turbulent mixer. Nonetheless, the similarity of scattering profiles with comparable LC phases within a probed timeframe and mixing conditions (e.g. 45-637 ms for yRNA LNPs at 3:1 FRR), provides support for the consistency of the observations.

The *p*(r) profiles obtained for eLNPs showed no systematic evolution in particle dimensions over the investigated time range, similar to the behavior observed for loaded LNPs. The corresponding R_g_ values remained broadly comparable from the earliest measurable time point to the final time point, indicating that no substantial change in the characteristic particle dimensions was resolved following initial particle formation. Both empty and loaded LNPs appear to attain their overall particle dimensions within the earliest accessible time point following turbulent mixing.

The data presented in **Figure 3** indicates that the primary particle population has already formed at the earliest probed residence time of 45 ms. The lowest residence time measured represents a physical limitation of the high-throughput MIVM-SAXS setup including effluent tubing connections. To experimentally access the shortest possible residence time for LNPs produced at a 3:1 FRR using the quartz capillary flow-cell, a novel mixing setup was constructed. Using a resin 3D printer, we fabricated a bespoke T-junction mixer with a 1 mm internal diameter (ID) and a threaded (1/4”-28) outlet channel that allowed the mixer to be directly fitted onto the flow-cell quartz capillary, to bring fluid residence time at the point of X-ray exposure down to 17.4 ms (**Figure 4b**). LNPs encapsulating polyA were produced at 3:1 FRR at a TFR of 140 mL/min for continuous nanoprecipitation using the fabricated T-mixer, and SAXS frames were acquired immediately downstream. Even at a timescale of 17.4 ms, the earliest accessible timepoint for 3:1 mixers in this work, the LNPs are fully formed, suggesting that the onset of assembly precedes the earliest residence time that can be experimentally probed at a 3:1 FRR, and that LNPs in general assemble rapidly during turbulent mixing conditions.

**Figure 4.**
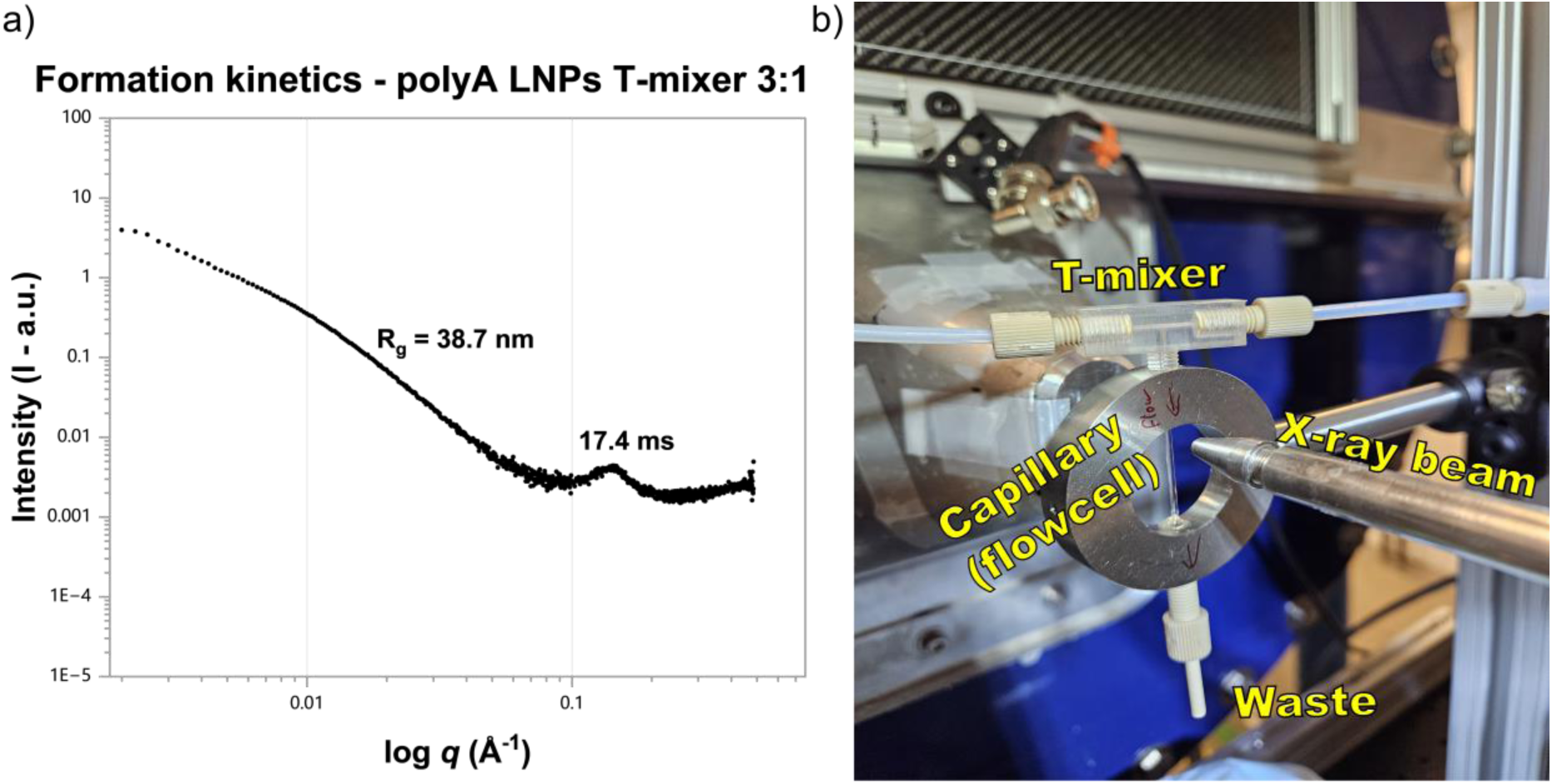
a) Flowthrough T-mixer SAXS measurements for *in situ* nanoprecipitation of polyA-loaded LNPs. Measurements were carried out at a fluid residence time of 17.4 ms post-mixing at a TFR of 140 mL/min and an FRR of 3:1 Aq:Org b) Labelled image of in-house fabricated T-mixer with ¼-28” threading which can be screwed into the capillary flow-cell, minimizing fluid residence time to 17.4 ms at the point of acquisition. The T-mixer was run at a TFR of 140 mL/min and an FRR of 3:1 Aq:Org.

### A flowrate ratio of 1:1 is associated with reduced order in the core of primary LNPs within milliseconds of formation

To determine whether the observed assembly kinetics were specific to the initial mixing and supersaturation conditions provided by a 3:1 FRR, we sought to probe LNP formation at a FRR of 1:1, where the solvent volume fraction was significantly higher. To this end, we applied an MIVM-SAXS and CIJM-SAXS setup (**Figure 1**) to investigate the formation kinetics of LNPs encapsulating yRNA produced at a 1:1 FRR and a TFR of 160 mL/min and 120 mL/min, respectively. **Figure 5a** shows the resulting scattering patterns for the MIVM 1:1 experiment, with comparable observations as the MIVM 3:1 results in **Figure 3**. LNPs produced at a 1:1 FRR using either the MIVM (**Figure 5a**) or the CIJM (**Figure 5c**) appeared to be fully formed by the probed timeframe of 32-251 ms. Due to the considerable difference in the fusogenicity of LNPs at 50% v/v of ethanol compared to 25% v/v, there is no size characterization information on 1:1 formulations prior to quenching (dilution post-mixing), as the particles would rapidly coalesce and ultimately aggregate in these conditions (within seconds). The setups employed in **Figure 1** allow for acquisition of SAXS patterns milliseconds after LNP formation and therefore provide a unique snapshot of the short-lived structural properties of LNPs formed at high solvent volume fractions. Interestingly, the LNPs produced at a 1:1 FRR exhibited a broader and more diffuse LC phase peak that was shifted towards lower-*q*, in contrast to the peaks observed in the 3:1 mixing experiments, which were more pronounced with a smaller *d*-spacing (**Figure 5b**). The breadth of these peaks reflects an altered domain size of the ordered phase within the LNP core and overall less uniform ordering..

**Figure 5.**
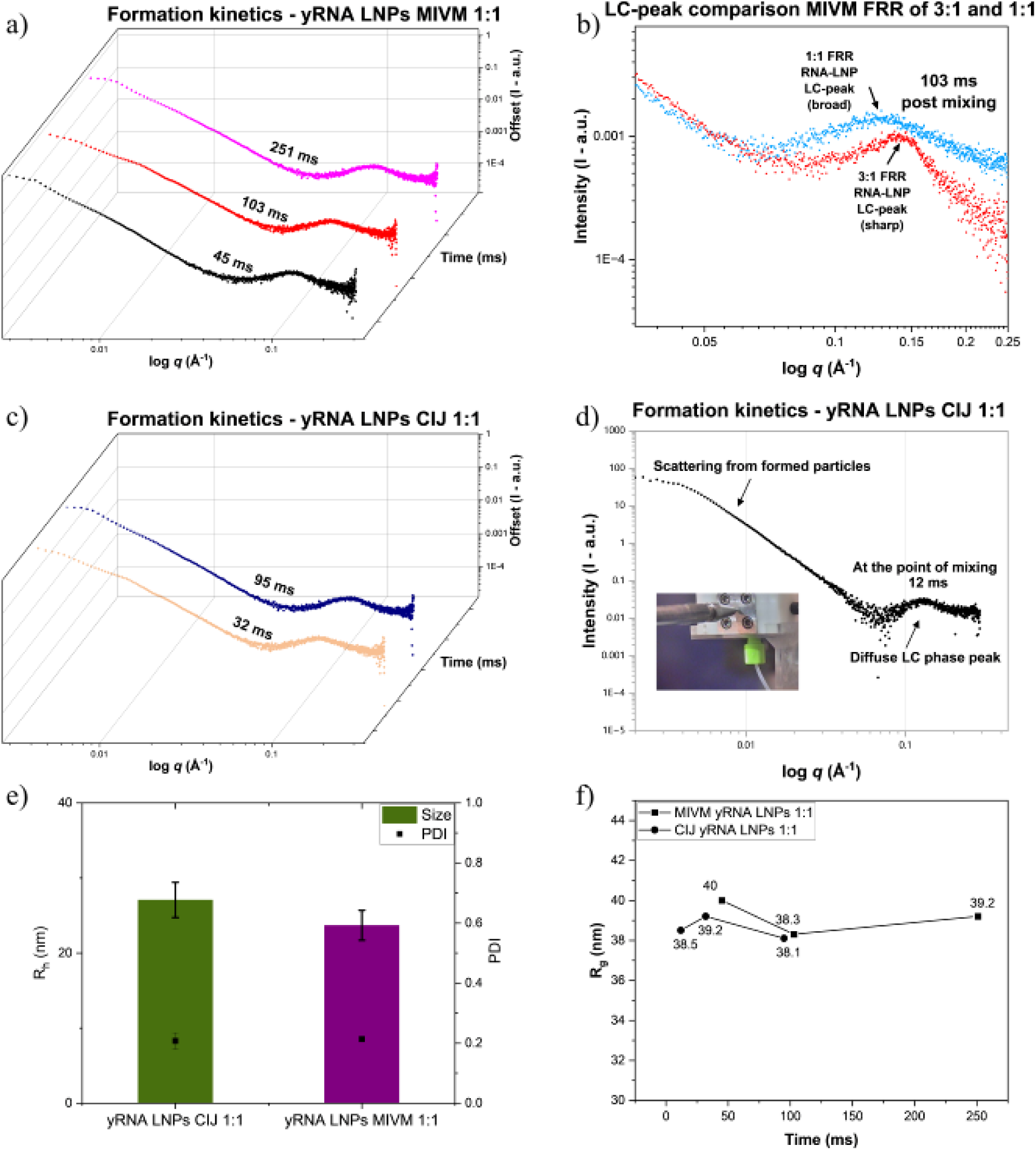
a) Flowthrough MIVM-SAXS measurements for *in situ* nanoprecipitation of yRNA loaded LNPs. Measurements were carried out at a fluid residence time of 45, 103, and 251 ms post-mixing at a TFR of 160 mL/min and an FRR of 1:1 Aq:Org. b) Comparison of LC-phase peak between yRNA-loaded LNPs formed at a FRR of 3:1 and 1:1 at 103 ms post-mixing in an MIVM. c) Flowthrough CIJM-SAXS measurements for *in situ* nanoprecipitation of yRNA-loaded LNPs. Measurements were carried out at a fluid residence time of 32 and 94 ms post-mixing at a TFR of 120 mL/min and an FRR of 1:1 Aq:Org. d) SAXS pattern obtained from scattering in a flowthrough CIJM setup with a quartz window, showing LNP formation at the point of mixing (12 ms). e) DLS measurements showing the intensity weighted z-average hydrodynamic radius R_h_ for the fully formed LNPs made using the MIVM and CIJM at a 1:1 FRR, quenched to 1% EtOH v/v (n = 3, ±SD). f) R_g_ (nm) extracted from each timepoint for yRNA-loaded LNPs formed at a 1:1 FRR plotted against fluid residence time (ms).

This additional disorder in the core of the LNPs produced at a 1:1 FRR, compared to 3:1, can be attributed to the increased fluidity of the lipids in the core, driven by a higher overall solubility in the medium and solvent uptake in or swelling of the lipid phases. This fluidity hinders the tight packing of the lipids and RNA and results in disordered domains. Importantly, SAXS analysis of fully processed LNPs prepared at a 3:1 FRR and a 1:1 FRR, with the 1:1 FRR immediately quenched, show comparable internal packing^10^. This suggests that the final relaxation of the internal ordering of LNPs produced at a 1:1 FRR is not dictated by primary mixing and the resulting self-assembly, but rather the downstream post-mixing processing that shifts the solvent quality towards a higher volume fraction of water, leading to lipid rearrangement and tighter packing, recovering the same high-*q* LC phase scattering signature. This was recently highlighted from a formulation control perspective, whereby timed quenching of a formulation produced at a 1:1 FRR, and therefore kinetic arrest of the effluent particles produced by an impinging jets mixer, resulted in varying LNP physicochemical and morphological properties^27^.

Importantly, the increased solvent volume fraction in the medium did not seem to impact overall particle formation kinetics. Using the bespoke engineered CIJM with a quartz window, SAXS patterns were acquired at the point of mixing (12 ms residence time, 120 mL/min). **Figure 5d** shows that LNP formation is rapid even in the presence of 50% EtOH v/v, as evident by particle scattering at low-*q* and the appearance of an LC-phase peak at high-*q*. The size of the particles (extracted using the *p*(r) function – plots in **Figure S1**) did not evolve over the probed timeframe of 12-251 ms and was within the PSD measured by DLS for the MIVM and CIJM (**Figure 5e and 5f**).

Overall, LNP self-assembly and initial structural maturation were concluded within milliseconds after rapid turbulent micromixing, as evidenced by the observed LNP particle scattering with internal ordering recorded at the earliest accessible time point of 12 ms and 17.4 ms (1:1 and 3:1 FRR, respectively) in our experimental setups, which still satisfies the constraint of Da < 1 for Flash Nanoprecipitation. This range for LNP formation kinetics shows good agreement with recently reported population balance equations that aimed to simulate the nucleation and growth kinetics of the resulting PSD of DODMA LNPs^5^. The model estimated stabilization of the PSD (not driven by delayed coalescence) to conclude within ∼60 ms, which is a longer timescale than what was observed experimentally for the average population of particles to show structural maturation as measurable by SAXS. The difference likely arises from the fact that PBE-based simulations describe the convergence of a PSD, rather than the onset of fully formed individual particles, that result in an electron density contrast measurable by SAXS^28^. Additionally, the PBE models do not hold information on internal LC phase formation, which is a central aspect of the final particle morphology of LNPs.

Notably, Yu *et al.* demonstrated that the precipitation and complexation kinetics of model LNP core material with polyA concluded within ∼20 ms^29^. This was measured as the AUC evolution of the associated structural Bragg peak that appears due to ionizable lipid-nucleic acid complexation. In that work, significant differences were observed when the size and nature of the cargo was varied, with mRNA and DNA showing slower complexation kinetics. Such differences were not observed in the current study, likely due to the structural similarity between polyA and yeast RNA but also the lower Da achieved with the Flash NanoPrecipitation regime turbulent mixers. Additionally, it is worth noting that the correlation peak associated with RNA internalization observed at the earliest timepoints in both **Figure 4** and **Figure 5**, was fully matured within 17.4 ms and 12 ms, compared to the slower assembly observed using the model core material (ionizable lipid with monoolein stabilizer) by Yu, *et al*. This discrepancy between model core systems and clinical multi-component LNPs is possibly explained by the role of cholesterol in particle formation. The cholesterol in the clinical LNP formulations is purely hydrophobic, compared to the amphiphilic nature of the remaining three components. The precipitation and subsequent aggregation of LNPs are therefore likely to be templated by cholesterol acting as a hydrophobic nucleus alongside the ionizable lipid, facilitating rapid RNA complexation.

Recently, Gilbert *et al* investigated the structural evolution of LNPs during microfluidic mixing. Using an X-ray-transparent staggered herringbone mixer, they were able to acquire frames at several points along the mixing channel^30^. Interestingly, low-*q* scattering could be detected along the mixing channel, described as arising from the formation of “protoLNPs” – essentially initial aggregates of only lipids and lipids with nucleic acids prior to structural maturation. Due to this, no LC-phase was detected in these *in-situ* studies of polyA-LNPs inside the microfluidic channels and only matured after mixing and downstream processing outside the mixer. Our current study, which used turbulent mixing expected to homogenize the solvent and antisolvent in under 3 ms, could not decouple particle formation from internal LC-phase formation even at the point of mixing, where an LC-phase peak was present. This is attributed to the rapid nature of particle formation under conditions dictated by DLA, triggered due to faster solvent-antisolvent mixing than particle assembly at low Da number. The formation of particles at Da > 1 is generally considered to be governed by mixing dynamics, and that local gradients of solvent and anti-solvent can form. Maeki *et al* showed that liposome size in microfluidic chaotic mixers is governed by the residence time of lipid intermediates within a critical ethanol concentration window^31^. While the final size of LNPs is comparable between microfluidic mixing and CIJM mixing, it is possible that due to these local gradients, RNA loaded LNPs may assemble through different pathways. Together, these findings suggest that LNP self-assembly occurs through distinct, mixing dependent regimes, which is discussed in the following section.

### Modelling mixing-dependent self-assembly behavior of RNA-LNPs

The following section puts forth a diffusion limited aggregation model to approximate the time scale of LNP assembly, assuming rapid micromixing from the turbulent flow mixes the solvent and antisolvent streams faster than LNP assembly (Da < 1). RNA may be encapsulated through *in situ* loading (ISL), contemporaneous with the LNP assembly or through *post hoc* loading (PHL) with RNA added to already fully formed eLNPs. The effects of laminar vs turbulent flow, i.e. mixing that is slower (Da > 1) or faster (Da < 1) than the ∼10 ms self-assembly time, are another consideration. There are therefore four regimes to consider: laminar vs. turbulent mixing for LNP assembly, and ISL vs. PHL for RNA cargo encapsulation. We have considered (and measured) two of these, PHL in the laminar vs. turbulent regimes, previously^32^, so this work will present new models for ISL in the two flow regimes. These suggest distinct RNA loading mechanisms under turbulent vs. laminar mixing. As an extension, we also consider RNA molecular weight as a parameter (siRNA, various kinds of mRNA, and DNA structures have orders of magnitude different sizes and encapsulation time scales), and we end with a strategy, grounded in the models for all four regimes and with supporting experimental data, for maximizing RNA loading homogeneity across LNPs in a population. A two-dimensional matrix illustrating the four regimes discussed is presented in **Figure 6**, emphasizing their distinct mixing, LNP formation, and RNA encapsulation pathways.

**Figure 6.**
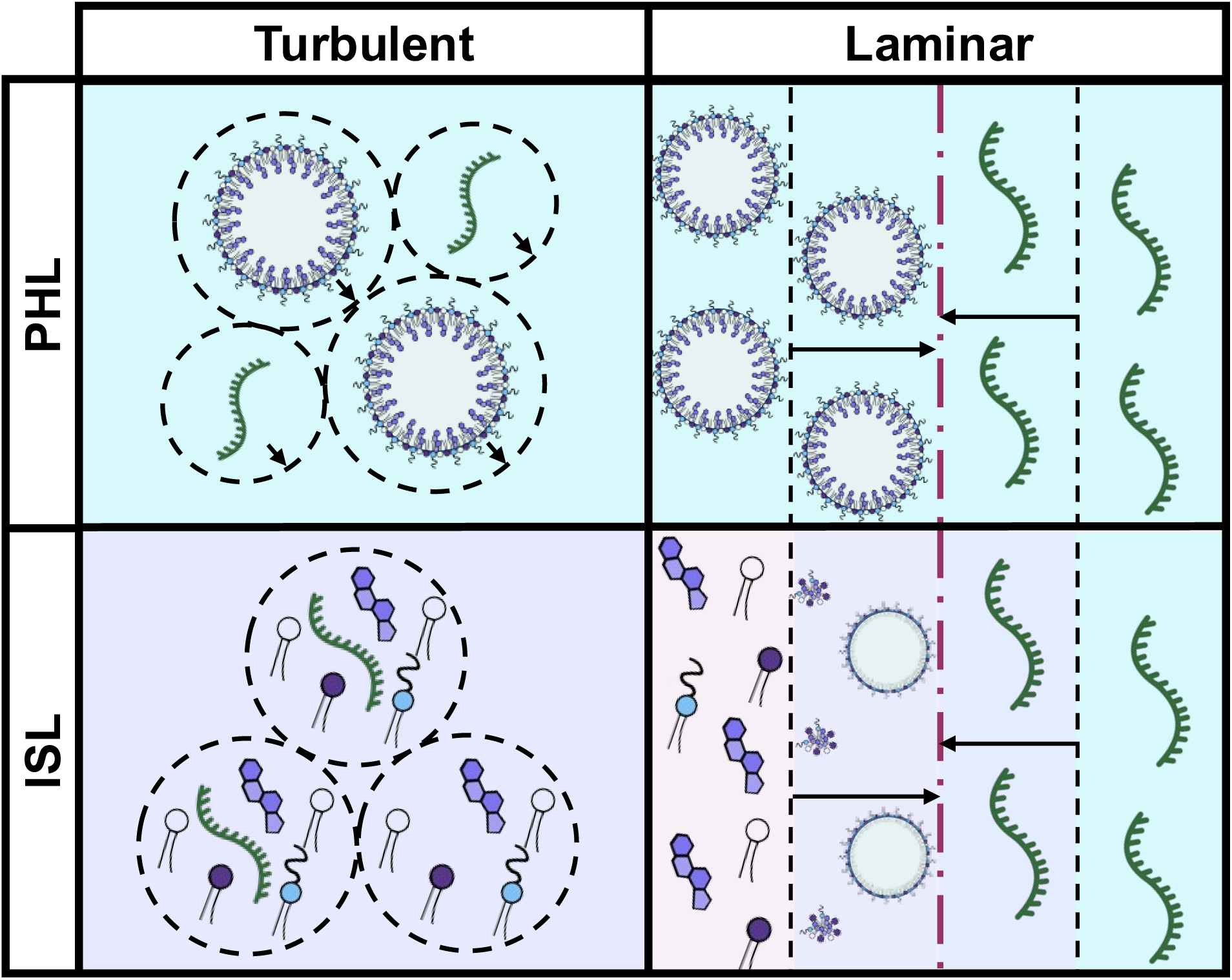
A conceptual two-dimensional matrix summarizing the proposed self-assembly regimes under conventional *in situ* loading (ISL) and *post hoc* loading (PHL) across laminar and turbulent mixing conditions. Turbulent mixing creates homogeneous supersaturated solutions, with nucleation and growth also occurring in an approximately homogeneous manner. Laminar mixing creates a well-defined interface between the two streams with solvent-antisolvent mixing occurring over a larger length scale and much longer time scale leading to heterogeneous supersaturation, nucleation, and growth of LNPs. Note that in the PHL case, both streams are aqueous for formed eLNPs mixed against RNA solution while in the ISL case, one stream is organic (ethanolic) and the other is aqueous. All aqueous streams contain buffer ions (10 mM, pH 5 acetic acid/sodium acetate buffer here).

### *Post hoc* loading dominates cargo internalization under laminar flow *in situ* formulation

LNP assembly under laminar mixing conditions was modeled as a lipid and RNA diffusion problem, where lipids must diffuse to a zone adjacent to the water-ethanol interface to encounter sufficient antisolvent mixing to undergo precipitation. RNA requires electrostatic interaction with the ionizable lipid for encapsulation, so any RNA cargo must also diffuse to the water-ethanol interface to find its lipid vehicle components. Water, ethanol, and lipids all diffuse much more rapidly than hydrodynamically large RNAs (mRNA, saRNA, *etc.*), while siRNAs are a special case which have a higher number density and diffusivity. We hypothesize that under laminar flow conditions, in which mixing is driven largely by diffusion over length scales defined by the dimensions of the mixing channels in the device (∼100µm) rather than features arising from the flow (in CIJ turbulent mixing, lamellae on the order of 1-5µm), there is sufficient time for water to diffuse far enough into the ethanol stream to induce supersaturation of lipids, triggering lipid self-assembly into vesicles, before RNA can diffuse from the aqueous stream to the growing lipid aggregate, especially for high MW RNAs. This would lead to the formation of eLNPs (or lipidic aggregates which will mature through further growth into eLNPs) that subsequently internalize RNA through what is fundamentally a *post hoc* loading rather than *in situ* co-precipitation, a mechanism not previously identified for these kinds of mixers. The mechanism is illustrated in the “Laminar ISL” panel in **Figure 6** where lipids approaching the mixed water-ethanol interface assemble into lipid particles before complexing with and encapsulating RNA.

In essence, this model encodes the differences in hydrodynamic size and diffusivity between lipids and RNA in the laminar flow mixer geometries into the relative timescales of solvent diffusion, lipid self-assembly, and RNA diffusion. Lipid self-assembly into eLNPs was modeled using a Smoluchowski-type step-growth model, which gave a characteristic self-assembly time of 1.29 ms for a 40 nm diameter eLNP with a 3 nm bilayer thickness (**Supporting Information**)^18^. This timescale is consistent with our SAXS experimental findings and with previous models in the literature^5^. It should be noted that this model assumes perfectly productive collisions between lipid molecules (a DLA mechanism), which may not occur at very short initial timescales before, in laminar flow, enough water has diffused to drive sufficient supersaturation. In any case, the model provides a useful, lower bound order of magnitude estimate for the assembly timescale window in which RNA arriving after this limit is encapsulated by an integral *post hoc* step to eLNPs rather than a true contemporaneous co-precipitation expected for *in situ* concept.

RNA diffusion was analyzed over the 1.29 ms eLNP assembly interval using a simplified laminar flow model in which the ethanol stream was discretized into strips 143 nm in width, which corresponds to the diameter of a spherical packet of ethanol solution that contains enough lipids to produce one eLNP (**Supporting Information**). We utilized prior measurements of RNA transcript hydrodynamic radii^32,33^ to compute the diffusivity of different molecular weights of RNA polymers, to determine the fraction of RNA molecules in the aqueous stream that could diffuse to the halfway point of this first strip (i.e. 71.7 nm into the strip, regardless of starting location, assuming the water layer is 3 times thicker than the ethanol layer and that RNAs are distributed evenly in this buffer stream) before lipid self-assembly was complete. Note that the enhanced colloidal properties seen from running laminar mixers at 3:1 aqueous:organic flow rate ratio worsens the RNA diffusion problem by diluting the already slow-diffusing RNA macromolecules^10,27^. For RNA molecular weights from 3.4 to 2000 kDa, fewer than 1% of molecules could diffuse to the halfway point within the 1.29 ms lipid self-assembly time (**Supporting Information**), and this percentage decreases even further for the subsequent strips of ethanol, proceeding further into the ethanol phase away from the water-ethanol mixing interface. Thus, only a small fraction of RNA could possibly encounter lipids to undergo *in situ* loading by complexing first with molecularly dissolved lipids before self-assembly of eLNPs is complete. The remaining RNA is predicted to interact with pre-formed eLNPs, resulting in *post hoc* loading as the primary mechanism of RNA internalization, in which RNA is encapsulated via RNA adsorption onto cationic lipid nanoparticles^34^. We have modeled RNA *post hoc* loading into eLNPs under laminar flow conditions previously and provided mechanistic explanations for heterogeneity in RNA distribution across the LNP population due to differences in particle diffusivity and initial spatial distribution.^32^ LNPs manufactured under laminar conditions with RNA present in the aqueous stream are labeled as *in situ* loading but proceed via an integral *post hoc* process due to the aggregation of lipids before most RNAs will ever encounter the lipid-rich phase.

### Formation of eLNPs is likely when high molecular weight cargo is encapsulated under turbulent flow *in situ* nanoprecipitation

A major consideration in the rapid mixing ISL case is the stoichiometry between cargo and vehicles. Consider perfect, instantaneous mixing of the ethanolic lipid stream and aqueous buffer RNA stream: the mixed system is a supersaturated solution of lipids (in the mixed water-ethanol solvent), with hydrodynamically large mRNA molecules distributed throughout this solution at very low number density compared to the lipids. At high mRNA molecular weight, every mRNA molecule will be surrounded by lipids and become encapsulated, but not every lipid will be near an mRNA, and clusters of such lipids will form into eLNPs. The result is a mixed population of ‘primary vesicles,’ a term we use to denote the non-fused, liposome-like LNPs formed at low pH, which are primarily electrostatically stabilized with sparse PEG-lipid on the surface and large, positive surface charge densities. When these primary vesicles are neutralized, by exchanging the acidic buffer for a neutral buffer, the ionizable lipid generated cationic surface potential vanishes. This loss of electrostatic stabilization initiates particle aggregation and fusion to the point where specific surface area decreases and PEG surface density increases enough to sterically stabilize the resulting LNPs. Approximately 14 vesicles fuse into 1 LNP (**Supporting Information**). If the population of primary vesicles is heterogeneous in RNA loading, the resulting LNPs will also be. Therefore, identifying the RNA MW at which this crossover occurs is important, and alternate strategies are needed to homogeneously load high MW RNAs. We present sequential *post hoc* loading as a viable strategy for creating nearly uniformly loaded LNPs even with large polyanion cargoes.

We first considered the simple molar stoichiometry between lipid components, the vesicle-like primary LNPs they assemble into at low pH, and the RNA cargo by calculating the ratio of total lipid vesicles to number of nucleic acid polymers as a function of RNA molecular weight. When the number of cargo molecules is lower than the number of vesicles, some of the primary vesicles must be empty (eLNPs). Assuming spherical, water-filled eLNPs 40 nm diameter and with a bilayer 3 nm thick^8^, ∼17,000 lipids are required to form a single vesicle (details provided in the **Supporting Information**). From this, using a lipid feed concentration of 12 mg/mL and a twofold dilution upon mixing in a CIJ, the number of vesicles in the system can be estimated. The ratio of RNA molecules to vesicles can then be calculated for a range of nucleic acid molecular weights, at a constant N/P ratio of 5. This N/P ratio contains the molar ratio of lipids to RNA and is a defining characteristic of a formulation. When the ratio of nucleic acid polymers to vesicles falls below 1, the cargo cannot occupy every vesicle, and even under instantaneous mixing, eLNPs will form. The model predicts this will occur above an RNA molecular weight of approximately 581 kDa. To put this into perspective, the mRNA encoding the Cas9 protein from *Streptococcus pyogenes* (SpCas9) is roughly 1140 kDa.^35^ Since many mRNAs of interest for gene editing applications are on the order of 1000 kDa or larger, this analysis suggests that turbulent flow *in situ* nanoprecipitation is expected to produce a substantial population of eLNPs when encapsulating large RNA cargos, and that this heterogeneity in loading is not expected for siRNAs encapsulated *in situ* using turbulent flow. This concept is visualized in the “Turbulent ISL” panel in **Figure 6**.

### *Post hoc* loading under rapid turbulent flow mixing maximizes the homogeneity of high molecular weight RNAs distributed across LNPs in a population

The fundamental challenge encountered when attempting to encapsulate high molecular weight mRNAs into LNPs via *in situ* loading is that even if mixing is rapid enough to enable LNP self-assembly by DLA (as is typically desirable in nanoprecipitation), the principles of diffusion dictate that small, fast-diffusing lipids will form eLNPs away from large, sparse, slow-diffusing mRNA. Faster mixing cannot improve this situation.

Importantly, a recent study by Wilson and Prud’homme developed rules for co-encapsulating (via diffusion-limited aggregation in Flash NanoPrecipitation) two species that are significantly different in size^36^. They showed that to achieve stoichiometric distribution of the two species across a population of particles, the initial diffusion limited aggregation timescale of the two species must be within a factor of ∼30 from one another. The diffusivity-timescale mismatch between large mRNA molecules and small lipids exceeds this limit. *Post hoc* loading emerges as a solution, however. The diffusivity and number density of the large mRNA molecules (hydrodynamic radius ∼10 - 15 nm) is much more closely matched with that of the pre-formed LNPs (radius ∼20nm for the vesicles or ∼40 nm for the fused LNPs) than with that of the lipid molecules.

We prepared a series of LNP formulation encapsulating poly(styrene sulfonic acid) (PSS) as an analogue to RNA which is available in monodisperse, widely spaced molecular weights (1,830 Da; 68,300 Da; 504,400 Da; 1,244,00 Da; and 2,210,000 Da). The Raman spectrum of PSS has a strong, unique signal at ∼1600 cm^-1^ allowing spectroscopic measurement of cargo within a lipid matrix. While PSS is not reflective of the secondary structures in RNA (Watson-Crick base pairing and π-stacking associations), it is still an anionic polymer which is available in monodisperse, well-defined molecular weights (1,830 Da; 68,300 Da; and 504,400 Da used here from monodisperse SEC/GPC standards) which enabled us to test our hypothesis around the encapsulation of increasingly large polyanions through *in situ* and *post hoc* loading strategies. PSS-loaded LNPs supports analysis of cargo distribution between individual LNPs, including the detection of eLNPs in a dispersion when hydrodynamically large polyanions are used in an ISL production scheme. Attempts to make LNPs with the 1,244,00 or 2,210,000 Da PSS yielded colloidally stable but significantly larger LNPs (> 100 nm diameter at pH 4, > 150 nm diameter at pH 7.4), strongly suggesting the large hydrodynamic size of these PSS coils were interacting with and condensing multiple LNPs. Further analysis utilized the 504,400 Da and smaller PSS to avoid this effect.

The molecular weight of the PSS cargo creates two distinct regimes in the SPARTA data for turbulent (CIJM), *in situ* loaded LNPs: for 1,830 or 68,300 Da PSS cargoes, particles show practically all LNPs contain PSS cargo with rare eLNPs (under 1% of measured particles), but for 504,400 Da (52.1% empty) cargo the dispersion contained more eLNPs and cargo-loaded LNPs became increasingly rare. Switching the loading strategy to *post hoc* loading in the same turbulent mixer delivered a striking result where effectively all (over 98%) LNPs contained PSS cargo, for all PSS molecular weights from 1,830 Da to 504,400 Da. Both the ISL and PHL cases analyzed by SPARTA are shown in **Figure 7**. This result corroborates the models for ISL and PHL under turbulent flow rapid mixing and remarkably captures the approximate MW cutoff identified for sub-stoichiometric loading. Note that we also attempted an analogous experiment under laminar conditions using a tesla micromixer structure (NanoGenerator, PreciGenome; San Jose, CA) but this mixer introduced a contaminant into the product LNP dispersions which produced strong Raman scattering at 1600 and 1725 cm^-1^ wavenumbers, independent of whether or not PSS was added (see additional Raman spectra – **Figure S3**). Particle size distributions for the PSS-loaded LNPs are given in the supporting information (**Figure S4**).

**Figure 7:**
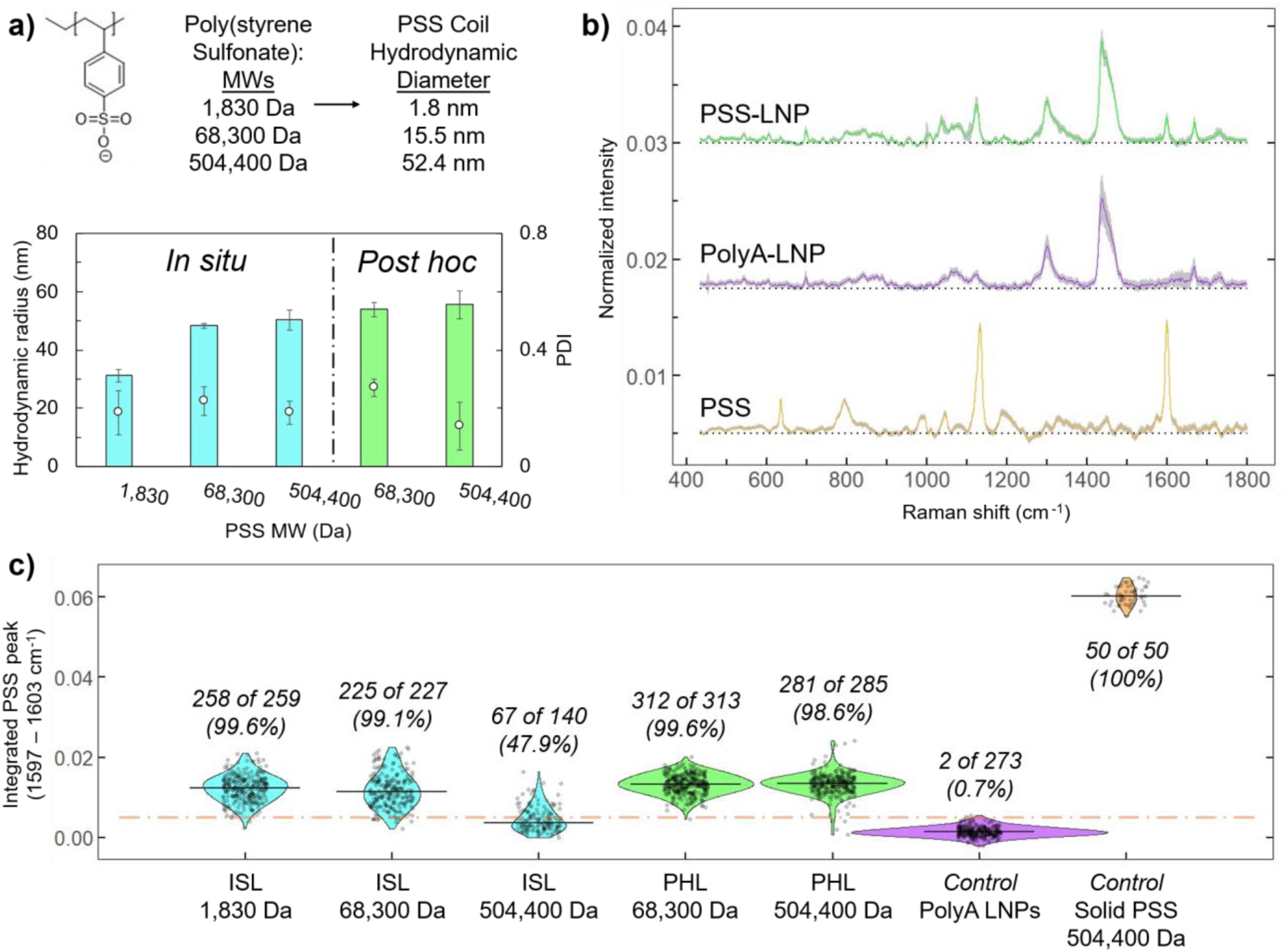
Single-particle Raman spectroscopy of PSS-laoded LNPs either *in situ* or *post hoc* under turbulent flow rapid mixing. a) poly(styrene sulfonate) (PSS) structure, molecular weights (MW), and coil hydrodynamic diameters (calculated from the PSS Mark-Houwink-Sakurada parameters) materials, with product LNP hydrodynamic radii (bars) and polydispersity indices (open circles) measured by DLS shown for *in situ* (ISL) and *post hoc* loading (PHL) strategies. b) Offset SPARTA normalized Raman spectra for solid PSS (bottom, orange line), polyA-loaded LNPs (middle, purple line), and PSS-loaded LNPs (top, green line). Zero-level for each spectrum is given by the dashed black lines. The standard deviation of the average intensity is shown as a gray shading around the line. c) Violin plot of PSS peak intensity for each LNP measured in the indicated combination of PSS MW and loading strategy. Annotated numbers indicate the number of PSS loaded particles of the total number of analyzed particles (with percentage loaded given). The PSS loading threshold indicated by the dashed orange line is defined as the 99^th^ percentile of the PSS-devoid polyA loaded LNP control^37^.

## Conclusion

This work provides experimental millisecond-resolved insight into LNP self-assembly kinetics using time-resolved SAXS coupled to high-throughput mixers, revealing that the primary structural features associated with LNP self-assembly occurs within the first milliseconds (under 12 ms) following turbulent mixing at both low q, describing overall particle diameter and shape, and also high q, describing particle internal structure due to lipid-RNA complex formation. Reducing the FRR from 3:1 to 1:1 resulted in distinct scattering profiles with similar low-*q* features but marked changes at high-*q* with broader LC-phase peaks, suggesting that the higher ethanol fraction (50% v/v) increased lipid fluidity during self-assembly, ultimately leading to a less ordered LNP nanostructure.

When compared to literature findings, these experimental results support a model in which LNP self-assembly proceeds through distinct, mixing-dependent assembly regimes for both the lipid aggregation into “empty” RNA-free LNPs and for encapsulation of the RNA payload in these lipid aggregates. We developed a conceptual framework describing four distinct LNP self-assembly regimes arising from the interplay between mixing mode (laminar vs turbulent) and loading strategy (*in* situ loading, ISL vs *post hoc* loading, PHL). We propose that the relative timescales of solvent-antisolvent mixing, and resulting lipid self-assembly, and RNA encapsulation through RNA-lipid complexation define the pathway by which LNPs are formed, providing a mechanistic basis for understanding how formulation and process conditions alter nanoparticle microstructure and population heterogeneity, ultimately influence formulation performance. These findings emphasize the importance of mixing rate and solvent quality on LNP assembly and RNA cargo encapsulation. Rapid micromixing and a deep solvent quench (more aqueous antisolvent) promote rapid formation of mature LNPs, which guides the design and operation of other post-mixing unit operations.

## Acknowledgements

This research was performed at beamline 12-ID-B on APS beam time awards (DOI: https://doi.org/10.46936/APS-189691/60013831 & https://doi.org/10.46936/APS-191740/60015470) from the Advanced Photon Source, a U.S. Department of Energy (DOE) Office of Science user facility operated for the DOE Office of Science by Argonne National Laboratory under Contract No. DE-AC02-06CH11357. We acknowledge the MAX IV Laboratory for beamtime on CoSAXS beamline under Proposal 20250632. Research conducted at MAX IV, a Swedish national user facility, is supported by Vetenskapsrådet (Swedish Research Council, VR) under contract 2018–07152, Vinnova (Swedish Governmental Agency for Innovation Systems) under contract 2018–04969, and Formas under contract 2019–02496. This research was supported by a grant from Eli Lilly and Company (USA). The TOC figure was partially made using biorender.com

## References

(1) Cullis, P. R.; Hope, M. J. Lipid Nanoparticle Systems for Enabling Gene Therapies. Molecular Therapy 2017, 25 (7), 1467–1475. 10.1016/j.ymthe.2017.03.013

(2) Brimacombe, C. A.; Kulkarni, J. A.; Cheng, M. H. Y.; An, K.; Witzigmann, D.; Cullis, P. R. Rational Design of Lipid Nanoparticles for Enabling Gene Therapies. Mol. Ther. Methods Clin. Dev. 2025, 33 (3), 101518. 10.1016/j.omtm.2025.101518

(3) Kulkarni, J. A.; Witzigmann, D.; Chen, S.; Cullis, P. R.; van der Meel, R. Lipid Nanoparticle Technology for Clinical Translation of siRNA Therapeutics. Acc. Chem. Res. 2019, 52 (9), 2435–2444. 10.1021/acs.accounts.9b00368

(4) Dasaro, S. R.; Johnson, L. D.; Salim, M.; He, V.; Amelemah, D.; Ponsonby-Thomas, E.; Barber, B.; Fong, W.-K.; Caukwell, J.; Peach, E.; Li, B.; Salvati Manni, L.; Kirby, N.; Warr, G. G.; Prud’homme, R. K.; Boyd, B. J.; Ristroph, K. Time-Resolved In Situ Small-Angle X-Ray Scattering to Determine the Kinetics of Formation of Liquid Crystalline Structure in the Core of Polymeric Nanoparticles during and after Turbulent Mixing. Nano Lett. 2025, 25 (17), 7075–7080. 10.1021/acs.nanolett.5c01095

(5) Shin, S.; Devos, C.; Udepurkar, A. P.; Inguva, P. K.; Myerson, A. S.; Braatz, R. D. Mechanistic Modeling of Lipid Nanoparticle (LNP) Precipitation via Population Balance Equations (PBEs). Chemical Engineering Journal 2025, 523, 167786. 10.1016/j.cej.2025.167786

(6) Leung, A. K. K.; Hafez, I. M.; Baoukina, S.; Belliveau, N. M.; Zhigaltsev, I. V; Afshinmanesh, E.; Tieleman, D. P.; Hansen, C. L.; Hope, M. J.; Cullis, P. R. Lipid Nanoparticles Containing siRNA Synthesized by Microfluidic Mixing Exhibit an Electron-Dense Nanostructured Core. The Journal of Physical Chemistry C 2012, 116 (34), 18440–18450. 10.1021/jp303267y

(7) Hald Albertsen, C.; Kulkarni, J. A.; Witzigmann, D.; Lind, M.; Petersson, K.; Simonsen, J. B. The Role of Lipid Components in Lipid Nanoparticles for Vaccines and Gene Therapy. Adv. Drug Deliv. Rev. 2022, 188, 114416. 10.1016/j.addr.2022.114416

(8) Kulkarni, J. A.; Witzigmann, D.; Leung, J.; van der Meel, R.; Zaifman, J.; Darjuan, M. M.; Grisch-Chan, H. M.; Thöny, B.; Tam, Y. Y. C.; Cullis, P. R. Fusion-Dependent Formation of Lipid Nanoparticles Containing Macromolecular Payloads. Nanoscale 2019, 11 (18), 9023–9031. 10.1039/C9NR02004G

(9) Mukherjee, S.; Shin, S.; Devos, C.; Udepurkar, A.; Sagmeister, P.; Trout, B. L.; Myerson, A. S.; Braatz, R. D. Understanding Size Distributions during Lipid Nanoparticle Manufacturing through Mechanistic Modeling. ACS Omega 2026, 11 (18), 26901–26908. 10.1021/acsomega.6c00100

(10) Bethiana, T.; Aljabbari, A.; Li, Y.; Mitra, H.; Baghbanbashi, M.; Harris, G. S.; Dasaro, S. R.; Masoomi, F.; Vago, F. S.; Hartzler, S. L.; Figueiredo, M.; Metskas, L. A.; Vlachos, P.; Ardekani, A. M.; Yeo, Y.; Ristroph, K. Effects of Different Mixing Techniques on mRNA Lipid Nanoparticle Physicochemistry and Biological Performance. Nat. Commun. 2026. 10.1038/s41467-026-72499-1

(11) Markwalter, C. E.; Pagels, R. F.; Wilson, B. K.; Ristroph, K. D.; Prud’homme, R. K. Flash NanoPrecipitation for the Encapsulation of Hydrophobic and Hydrophilic Compounds in Polymeric Nanoparticles. JoVE 2019, No. 143, e58757. doi:10.3791/58757

(12) Jensen, A. B.; Christensen, T. E. K.; Weninger, C.; Birkedal, H. Very Large-Scale Diffraction Investigations Enabled by a Matrix-Multiplication Facilitated Radial and Azimuthal Integration Algorithm: {\it MatFRAIA}. J. Synchrotron Radiat. 2022, 29 (6), 1420–1428. 10.1107/S1600577522008232

(13) Hopkins, J. B. BioXTAS RAW 2: New Developments for a Free Open-Source Program for Small-Angle Scattering Data Reduction and Analysis. J. Appl. Crystallogr. 2024, 57 (1), 194–208. 10.1107/S1600576723011019

(14) Semenyuk, A. V; Svergun, D. I. GNOM–a Program Package for Small-Angle Scattering Data Processing. J. Appl. Crystallogr. 1991, 24 (5), 537–540. 10.1107/S002188989100081X

(15) Johnson, B. K.; Prud’homme, R. K. Chemical Processing and Micromixing in Confined Impinging Jets. AIChE Journal 2003, 49 (9), 2264–2282. 10.1002/aic.690490905

(16) Subraveti, S. N.; Wilson, B. K.; Bizmark, N.; Liu, J.; Prud’homme, R. K. Synthesizing Lipid Nanoparticles by Turbulent Flow in Confined Impinging Jet Mixers. JoVE 2024, No. 210, e67047. doi:10.3791/67047

(17) Morozova, T. I.; Nikoubashman, A. Coil–Globule Collapse of Polystyrene Chains in Tetrahydrofuran–Water Mixtures. J. Phys. Chem. B 2018, 122 (7), 2130–2137. 10.1021/acs.jpcb.7b10603

(18) Pagels, R. F.; Edelstein, J.; Tang, C.; Prud’homme, R. K. Controlling and Predicting Nanoparticle Formation by Block Copolymer Directed Rapid Precipitations. Nano Lett. 2018, 18 (2), 1139–1144. 10.1021/acs.nanolett.7b04674

(19) Aburai, K.; Hatanaka, K.; Takano, S.; Fujii, S.; Sakurai, K. Characterizing an siRNA-Containing Lipid-Nanoparticle Prepared by a Microfluidic Reactor: Small-Angle X-Ray Scattering and Cryotransmission Electron Microscopic Studies. Langmuir 2020, 36 (42), 12545–12554. 10.1021/acs.langmuir.0c01079

(20) Padilla, M. S.; Shepherd, S. J.; Hanna, A. R.; Kurnik, M.; Zhang, X.; Chen, M.; Byrnes, J.; Joseph, R. A.; Yamagata, H. M.; Ricciardi, A. S.; Mrksich, K.; Issadore, D.; Gupta, K.; Mitchell, M. J. Elucidating Lipid Nanoparticle Properties and Structure through Biophysical Analyses. Nat. Biotechnol. 2025. 10.1038/s41587-025-02855-x

(21) Kulkarni, J. A.; Darjuan, M. M.; Mercer, J. E.; Chen, S.; van der Meel, R.; Thewalt, J. L.; Tam, Y. Y. C.; Cullis, P. R. On the Formation and Morphology of Lipid Nanoparticles Containing Ionizable Cationic Lipids and siRNA. ACS Nano 2018, 12 (5), 4787–4795. 10.1021/acsnano.8b01516

(22) Li, S.; Hu, Y.; Lin, J.; Schneiderman, Z.; Shao, F.; Wei, L.; Li, A.; Hsieh, K.; Kokkoli, E.; Curk, T.; Mao, H.-Q.; Wang, T.-H. Single-Particle Spectroscopic Chromatography Reveals Heterogeneous RNA Loading and Size Correlations in Lipid Nanoparticles. ACS Nano 2024, 18 (24), 15729–15743. 10.1021/acsnano.4c02341

(23) Li, S.; Hu, Y.; Li, A.; Lin, J.; Hsieh, K.; Schneiderman, Z.; Zhang, P.; Zhu, Y.; Qiu, C.; Kokkoli, E.; Wang, T.-H.; Mao, H.-Q. Payload Distribution and Capacity of mRNA Lipid Nanoparticles. Nat. Commun. 2022, 13 (1), 5561. 10.1038/s41467-022-33157-4

(24) Li, S.; Hu, Y.; Li, A.; Lin, J.; Hsieh, K.; Schneiderman, Z.; Zhang, P.; Zhu, Y.; Qiu, C.; Kokkoli, E.; Wang, T.-H.; Mao, H.-Q. Payload Distribution and Capacity of mRNA Lipid Nanoparticles. Nat. Commun. 2022, 13 (1), 5561. 10.1038/s41467-022-33157-4

(25) Kolb, B.; Graewert, M.; Drexel, R.; Meier, F.; Raab, J.; Wilhelmy, C.; Nawroth, T.; Soloviov, D.; Haas, H.; Langguth, P. Advanced Quality and Comparability Assessment of mRNA-Loaded Lipid Nanoparticles: Absolute Size Distribution Profiles and Structure from AF4-Coupled Light and X-Ray Scattering Measurements. Anal. Chem. 2026, 98 (7), 5271–5283. 10.1021/acs.analchem.5c05911

(26) Yanez Arteta, M.; Kjellman, T.; Bartesaghi, S.; Wallin, S.; Wu, X.; Kvist, A. J.; Dabkowska, A.; Székely, N.; Radulescu, A.; Bergenholtz, J.; Lindfors, L. Successful Reprogramming of Cellular Protein Production through mRNA Delivered by Functionalized Lipid Nanoparticles. Proceedings of the National Academy of Sciences 2018, 115 (15), E3351–E3360. 10.1073/pnas.1720542115

(27) Devos, C.; Udepurkar, A.; Sagmeister, P.; Hodlewsky, A. S.; Chen, J.; Hatas, A.; Ostrovsky, N.; Al-Jazrawe, M.; Ren, J. I.; Liu, A. Y.; Braatz, R. D.; Myerson, A. S. Manufacturing mRNA-Loaded Lipid Nanoparticles with Precise Size and Morphology Control. ACS Nano 2025, 19 (38), 33991–34002. 10.1021/acsnano.5c09800

(28) Schwarzer, H.-C.; Schwertfirm, F.; Manhart, M.; Schmid, H.-J.; Peukert, W. Predictive Simulation of Nanoparticle Precipitation Based on the Population Balance Equation. Chem. Eng. Sci. 2006, 61 (1), 167–181. 10.1016/j.ces.2004.11.064

(29) Yu, H.; Angelova, A.; Angelov, B.; Dyett, B.; Matthews, L.; Zhang, Y.; El Mohamad, M.; Cai, X.; Valimehr, S.; Drummond, C. J.; Zhai, J. Real-Time pH-Dependent Self-Assembly of Ionisable Lipids from COVID-19 Vaccines and In Situ Nucleic Acid Complexation. Angew. Chem. Int. Ed. 2023, 62 (35), e202304977. 10.1002/anie.202304977

(30) Gilbert, J.; Sebastiani, F.; Arteta, M. Y.; Terry, A.; Fornell, A.; Russell, R.; Mahmoudi, N.; Nylander, T. Evolution of the Structure of Lipid Nanoparticles for Nucleic Acid Delivery: From in Situ Studies of Formulation to Colloidal Stability. J. Colloid Interface Sci. 2024, 660, 66–76. 10.1016/j.jcis.2023.12.165

(31) Maeki, M.; Fujishima, Y.; Sato, Y.; Yasui, T.; Kaji, N.; Ishida, A.; Tani, H.; Baba, Y.; Harashima, H.; Tokeshi, M. Understanding the Formation Mechanism of Lipid Nanoparticles in Microfluidic Devices with Chaotic Micromixers. PLoS One 2017, 12 (11), e0187962.

(32) Aljabbari, A.; Binion, H.; Dasaro, S. R.; Mitra, H.; Bethiana, T.; Harris, G.; Zuo, X.; Baghbanbashi, M.; Barrio-Zhang, A.; Herrera, D. P.; Figueiredo, M.; Wilson, B. K.; Ardekani, A.; Ristroph, K. RNA Post Hoc Loading into Empty Lipid Nanoparticles Occurs on Millisecond Timescales during Turbulent Mixing. bioRxiv 2026. 10.64898/2026.06.26.732967

(33) Borodavka, A.; Singaram, S. W.; Stockley, P. G.; Gelbart, W. M.; Ben-Shaul, A.; Tuma, R. Sizes of Long RNA Molecules Are Determined by the Branching Patterns of Their Secondary Structures. Biophys. J. 2016, 111 (10), 2077–2085. 10.1016/j.bpj.2016.10.014

(34) Kulkarni, J. A.; Thomson, S. B.; Zaifman, J.; Leung, J.; Wagner, P. K.; Hill, A.; Tam, Y. Y. C.; Cullis, P. R.; Petkau, T. L.; Leavitt, B. R. Spontaneous, Solvent-Free Entrapment of siRNA within Lipid Nanoparticles. Nanoscale 2020, 12 (47), 23959–23966. 10.1039/d0nr06816k

(35) Novus Biologicals. Recombinant S. pyogenes CRISPR-associated Protein 9, CF (9957-C9). https://www.novusbio.com/products/recombinant-s-pyogenes-crispr-associated-protein-9-cf_9957-c9 (accessed 2026-07-06)

(36) Wilson, B. K.; Prud’homme, R. K. Co-Encapsulation of Organic Polymers and Inorganic Superparamagnetic Iron Oxide Colloidal Crystals Requires Matched Diffusion Time Scales. Soft Matter 2024, 20 (41), 8312–8325. 10.1039/d4sm00935e

(37) Knaul, J. Z.; Bui, V. T.; Creber, K. A. M. Viscometric Constants for Sodium Polystyrene Sulfonate Standards in Solvents of 0.1 M CH3COOH/0.2 M NaCl and 0.2 M CH3COOH/0.1 M CH3COONa at 25.0 °C. J. Chem. Eng. Data 2000, 45 (3), 508–511. 10.1021/je990332n

