## Supporting information for "On the formation of lipid nanoparticles and encapsulation of RNA: kinetics, dynamics, and self-assembly regimes"

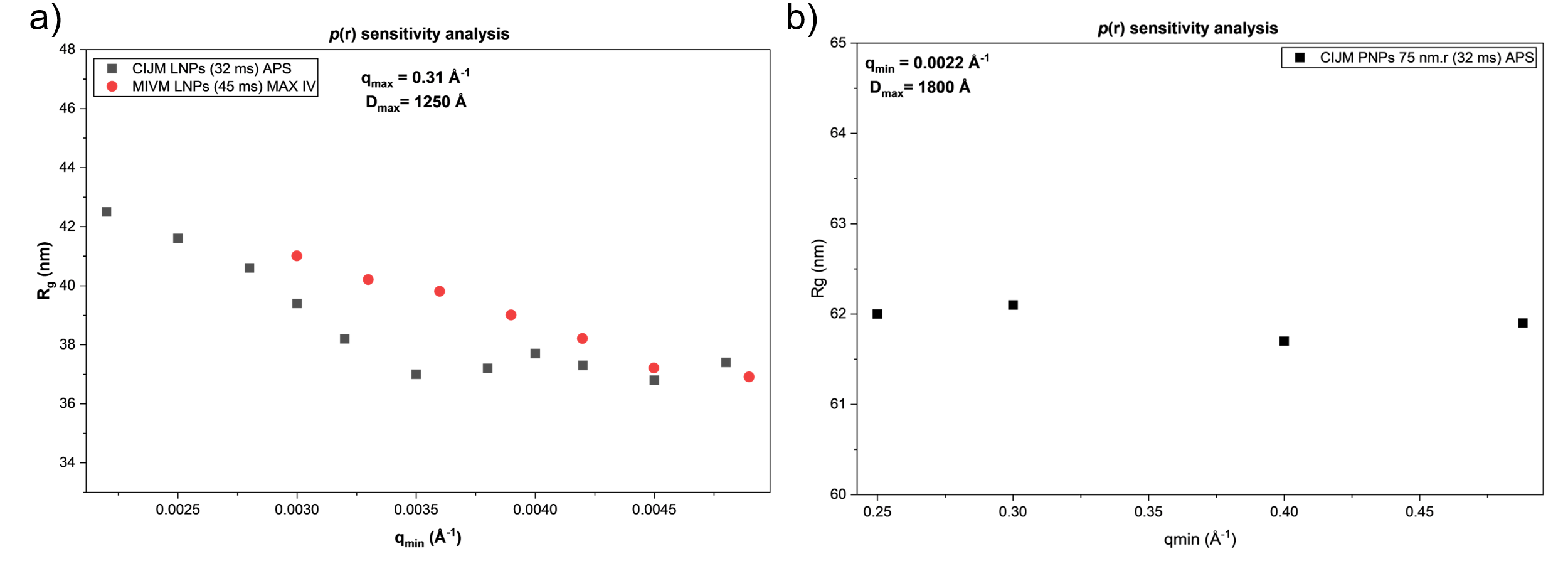

**Figure S1.** a) Sensitivity analysis for R_g_ extracted through *p*(r) analysis for the averaged scattering patterns of the flowthrough CIJM-SAXS and MIVM-SAXS measurements of yRNA-loaded LNPs at an FRR of 1:1 and a TFR of 120 mL/min. Extracted R_g_ at a fixed D_max_ and q_max_ is plotted against increasing q_min_. b) Sensitivity analysis for R_g_ extracted through *p*(r) analysis for the averaged scattering pattern of the flowthrough CIJM-SAXS and MIVM-SAXS measurements of large PNPs (75 nm.r) at an FRR of 1:1 and a TFR of 120 mL/min. Extracted R_g_ at a fixed D_max_ and q_min_ is plotted against increasing q_max_.

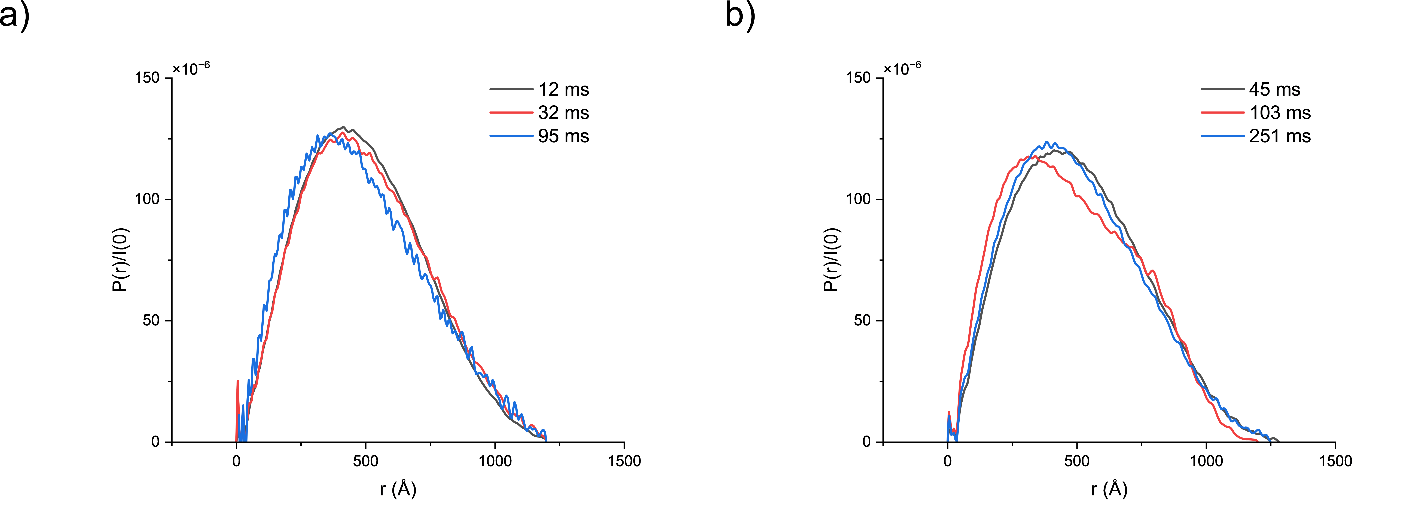

**Figure S2.** a) *p*(r) profiles for each averaged scattering pattern of the flowthrough CIJM-SAXS measurements of yRNA-loaded LNPs at 12, 32, and 95 ms, at an FRR of 1:1 and a TFR of 120 mL/min b) *p*(r) profiles for each averaged scattering pattern of the flowthrough MIVM-SAXS measurements of yRNA-loaded LNPs at 45, 103, and 251 ms, at an FRR of 1:1 and a TFR of 160 mL/min.

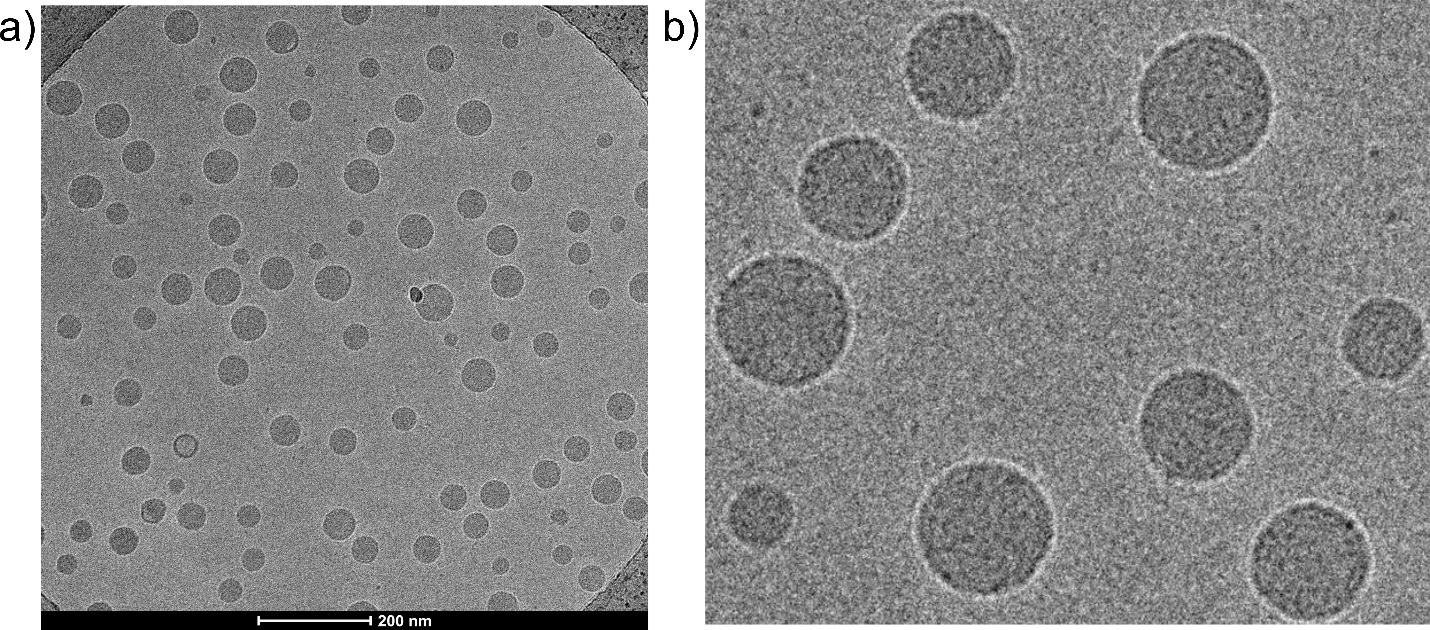

**Figure S3.** a) Cryo-TEM image showing LNPs produced at pH 5.0 (10 mM, acetate buffer) and dialyzed against pH 5.0 (10 mM, acetate buffer). b) Magnified cryo-TEM image from panel a, highlighting the condensed internal core of the LNPs at pH 5.0, similar to that observed for LNPs at pH 7.4, suggesting a broad neutralization of the ionizable lipid.

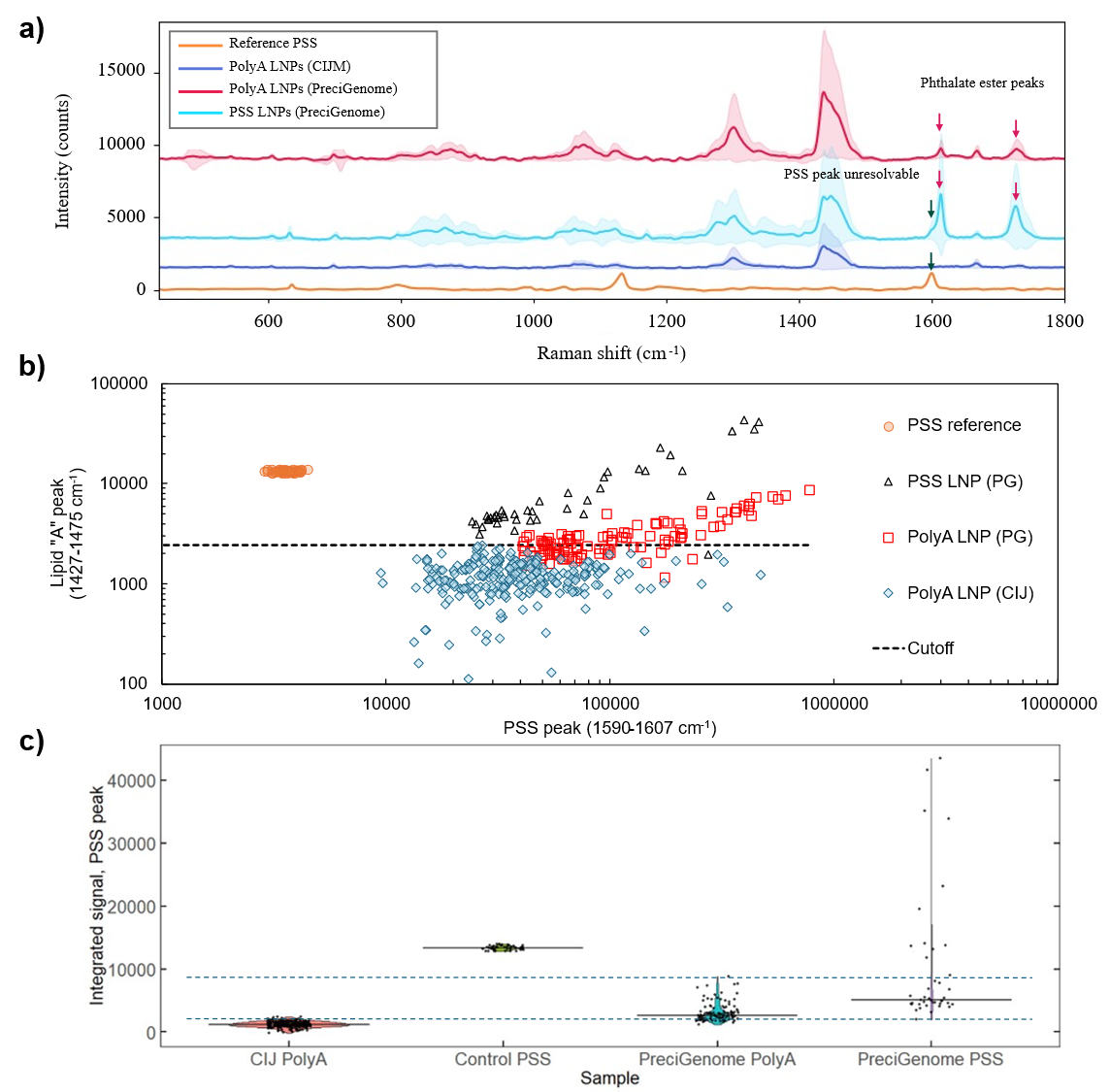

**Figure S4.** Single-particle Raman spectroscopy of LNPs loaded with either 504,400 Da PSS or polyA *in situ* using a tesla microstructure mixer (NanoGenerator, Precigenome, San Jose, California) or a CIJ. a) PolyA loaded LNPs made with Precigenome display unexpected additional peaks at 1600 cm^-1^ and 1725 cm^-1^ compared to polyA loaded CIJ LNPs, obscuring PSS signal and preventing analysis of PSS loading heterogeneity. b) Bivariate scatter of the lipid peak (range 1427 – 1475 cm^-1^ covers the “A” peak where all lipids have Raman scattering signal) against the PSS peak. The polyA loaded particles show signal comparable to PSS-loaded LNPs due to the exogenous component from the PreciGenome mixer. c) Violin plot of PSS peak data showing the polyA LNP from the PreciGenome PSS signal significantly spans the gap between polyA LNPs and the PSS standard (dashed lines showing span of PSS peak in the PreciGenome polyA LNPs) which overlaps with most particles measured for the PreciGenome PSS LNPs.

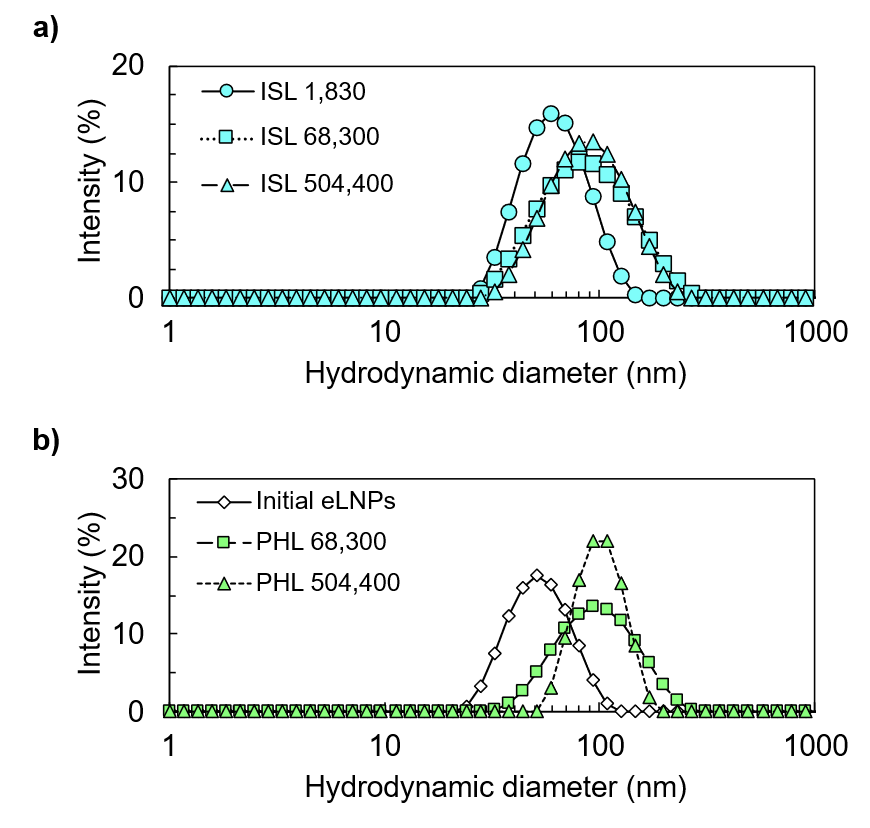

**Figure S5.** Particle size distribution for PSS-loaded LNPs characterized by SPARTA. a) *In situ* loaded LNPs with PSS 1,830; 68,300; and 504,400 Da molecular weights. b) *Post hoc* loaded LNPs showing the initial empty LNPs (at pH 4) and the dialyzed, PSS-loaded LNPs used for SPARTA measurements.

**Modeling calculations**

*Bilayer volume calculation*

To estimate the number of lipids required to form an empty vesicle of 40 nm diameter and 3 nm bilayer thickness (L_B_), the volume of the lipid bilayer must be determined. The bilayer volume can be found by subtracting the volume of the empty core of the vesicle from the entire spherical shell as presented in **Figure S3**. The outer radius, R_outer­_, is equal to 20 nm and the inner radius, R_inner_, is equal to R_outer_ – L_B_ = 17 nm. Using the volume of a sphere and given values of R_inner_ and R_outer_, the volume of the bilayer is determined by subtracting the inner volume from the outer volume. With the given dimensions, the bilayer volume is found to be ~1.29 x 10^4^ nm^3^.

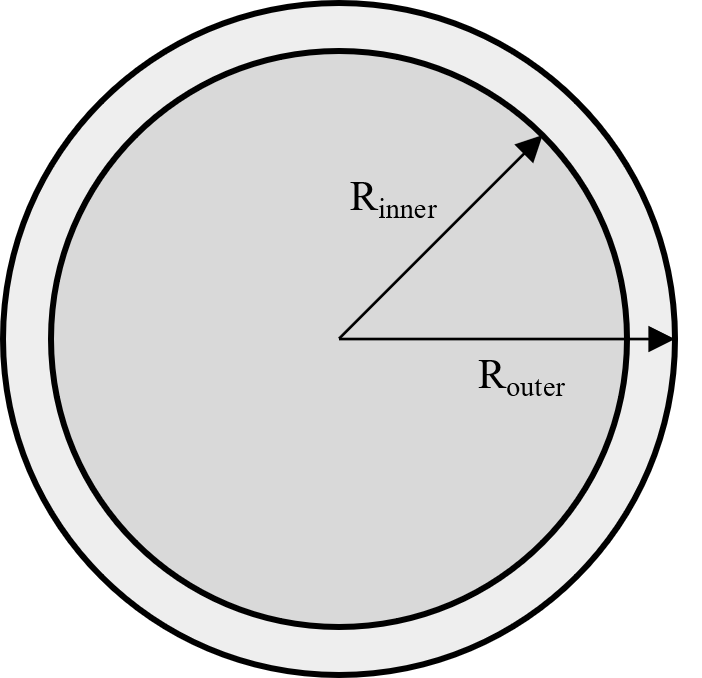

**Figure S6**. Bilayer volume determined by subtracting the inner sphere volume from the outer sphere volume, assuming a bilayer thickness (L_B_) of 3 nm.

*Lipids per vesicle calculation*

The number of individual lipid molecules in an LNP can be calculated from the measured geometry of the LNP (e.g., size, morphology) and the known composition of the lipid feed used. Individual lipid volumes (**Table S1)** were estimated using the molecular weight and density of lipids (molecular weight divided by the product of lipid density and Avogadro’s number yields units of volume):

**Table S1.** Average lipid volumes calculated through the weighted average of individual lipid volumes

| **Lipid** | **MW (g/mol)** | **ρ (g/mL)** | **Volume (nm^3^)** | **Molar Ratios** | **Weighted Volume (nm^3^)** | |
| --- | --- | --- | --- | --- | --- | --- |
| ALC-0315 | 766.29 | 0.90 | 1.41 | 0.500 | | 0.71 |
| DSPC | 790.14 | 1.00 | 1.31 | 0.100 | | 0.13 |
| Cholesterol | 386.65 | 1.02 | 0.63 | 0.385 | | 0.24 |
| DMG-PEG | 2509.2 | 1.00 | 4.17 x 0.2 = 0.83 | 0.015 | | 0.01 |
|  |  |  |  | Average (nm^3^/molecule): | | 1.09 |

Volumes were weighted by the molar ratio of lipids used and were summed to produce an average lipid volume of 1.09 nm^3^ per molecule. Only the lipidic portion of DMG-PEG2000 is assumed to contribute to the bilayer volume, thus a factor of 0.2 is used to adjust the volume that PEGylated lipid will occupy. Dividing the total bilayer volume by the average lipid volume per molecule gives n_lipids_ = 11,828 lipids necessary to form a singular eLNP.

*Number of lipid vesicles in mixing chamber*

Using the initial feed of composition of lipids (C_­lipids, feed_ = 12 mg/mL) and given that the lipid concentration is halved upon mixing with the aqueous feed, the number of vesicles can be calculated. This first requires knowing the mass of lipids (m_lipids_) that exist in the feed (**Eqn. E1**).

$$m_{Lipids}= \frac{n{}_{lipids}}{N_{A}}\times M_{W, avg}\times1000$$

(E1)

Where N_A­_ is Avogadro’s number (N_A_ = 6.022 x 10^23^ molecules/mole), and M_W, avg_ is the average molecular weight for lipids following the Onpattro molar ratio (~649 g/mol, **Table S2**).

**Table S2.** Average lipid molecular weight using lipid molecular weight and relative molar ratio of each lipid component.

| **Lipid** | **MW (g/mol)** | **Molar ratios** | **Weighted MW (Da)** |
| --- | --- | --- | --- |
| ALC-0315 | 766.29 | 0.500 | 383.15 |
| DSPC | 790.14 | 0.100 | 79.01 |
| Cholesterol | 386.65 | 0.385 | 148.86 |
| DMG-PEG | 2509.2 | 0.015 | 37.64 |
|  |  | Average MW (Da): | 648.66 |

The mixing chamber lipid concentration (C_lipids,chamber_) is 6 mg/mL. With a fixed mixing chamber volume (V_chamber_ = 0.1 mL), **Eqn. E2** can be used to determine the number of vesicle-forming regions (‘vesicles’ for simplicity; some of these will include an mRNA, and others will not) (n_vesicles_) that exist in the mixing chamber:

$$n_{vesicles}= \frac{C_{Lipids, chamber}}{m_{Lipids}}\times V_{Chamber}$$

(E2)

*Number of RNA in mixing chamber volume at a fixed N/P ratio:*

To determine how homogeneously distributed RNA cargo is in the mixer compared to assembly spheres, the number of RNA molecules at a fixed N/P ratio for a large range of molecular weights was calculated. Starting with the number of nucleotides of RNA, the molecular weight of a given RNA (M_w, RNA_) is found to be the number of nucleotides times the average nucleotide molecular weight (M_w, nucleotides_ ~ 340 g/mol). Using an N/P ratio of 5 and ALC-0315 as the ionizable lipid, the mass concentration of ALC-0315 (C_ALC-0315_ = 7.08 mg/mL) is used to determine the mass concentration of RNA in the feed (C_RNA, feed_). This can be seen in **Eqn. E3**:

$$C_{RNA,feed}=\left( \frac{C_{ALC-0315}}{5M_{W, ALC-0315}} \right)\times M_{W, Nucleotides}$$

(E3)

Where the molecular weight of ALC-0315 (M_w, ALC-0315_) is 766.29 g/mol.

The number of RNA molecules (n_RNA_) is then determined using **Eqn. E4**:

$$n_{RNA}=\left( \frac{C_{RNA, feed}}{M_{W, RNA}} \right)\times\left( \frac{N_{A}}{2} \right)\times\left( \frac{V_{Chamber}}{1000} \right)$$

(E4)

For the range of 3.4 kDa to 2040 kDa RNA at this fixed N/P ratio, the number of RNA molecules compared to assembly spheres was calculated. This is done by taking n_RNA_ divided by n_vesicles_. Initial calculations show this ratio of RNA/vesicles becoming less than 1 after ~1700 nucleotides or ~580 kDa as shown in **Figure S5**.

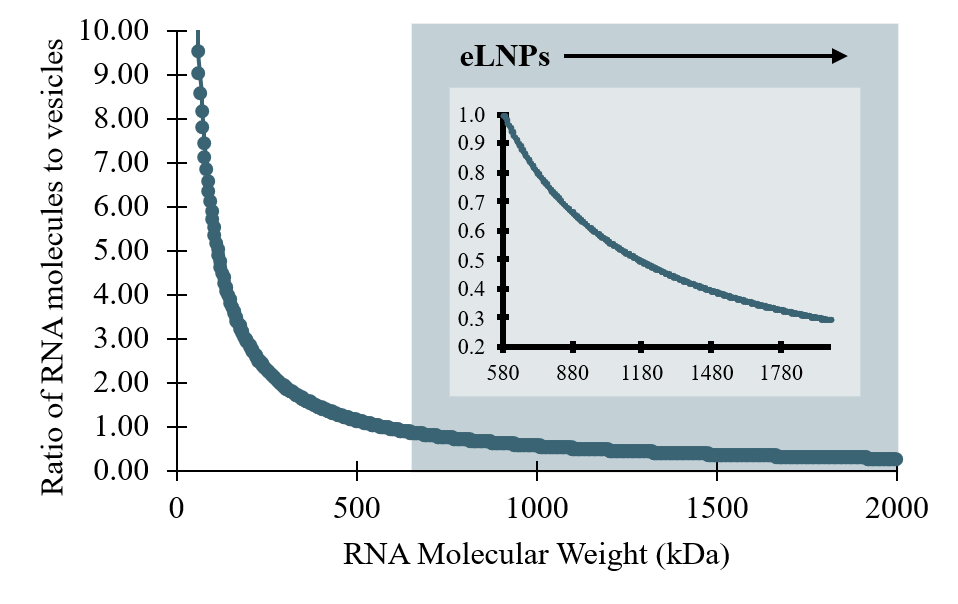

**Figure S5.** The ratio of RNA molecules to vesicles as a function of RNA molecular weight (kDa). This RNA to vesicle ratio decreases below unity at an RNA molecular weight of ~580 kDa.

*Timescale of lipid self-assembly*:

The kinetics of lipid self-assembly can be approximated using a Smoluchowski-type step-growth model. Starting from individual lipids, the number of doubling steps required to reach a particle containing ~12,000 lipids are calculated to be 13.53 (14 rounding up) using **Eqn. E5**:

$$Doubling steps=\log_{2} \left( \frac{N}{N_{o}} \right), N=11,863, N_{o}=1$$

(E5)

At each step, the number concentration of lipid aggregates is given by **Eqn. E6**:

$$C_{n}=\frac{c_{core}N_{A}}{M_{W, avg}n}$$

(E6)

Eqn. E6 gives the number concentration of lipids as a function of the core material concentration (C_n_ = 12 mg/mL), Avogadro’s number, the molecular weight of the core material (M_w, avg_), and the number of molecules composing the particle within a given growth step (n). The radius of aggregated lipids is then found through doubling of spheres.

Having the radius of aggregated lipids allows for the diffusivity of components to be determined via the Stokes-Einstein equation (**Eqn. E7**) where the diffusivity of each lipid (D_n_­) depends on the Boltzmann constant (k_B_ = 1.38 x 10^-23­^ m^2^kg/s^2^K**),** absolute temperature (T = 298.15 K), average solvent viscosity for a 25% ethanol in water composition (η = 0.0024 PaS)^1^, and the hydrodynamic radius of the primary lipid cluster (R_n_).

$$D_{n}=\frac{k{}_{B}T}{6\pi\eta R_{n}}$$

(E7)

Finally, with the number concentration of lipids, the size of the growing aggregate, and the diffusivity of the growing particle cluster, the time (in seconds) it takes to double in size can be determined in **Eqn. E8**:

$$\tau_{n\to2n}=\frac{1}{8\pi R_{n}D_{n}C_{n}}, \tau_{total}=\sum_{n=1}^{14} \tau_{n\to2n}$$

(E8)

Assuming 14 doubling steps are required and summing the time it takes for lipids to double, this process works out to be 1.29 ms for complete lipid self-assembly. This can be visualized in **Figure S6**.

| τ (ms) | Doubling step |
| --- | --- |
| 3.92638e-05 | 0 |
| 0.000117791 | 1 |
| 0.000274846 | 2 |
| 0.000588956 | 3 |
| 0.00121718 | 4 |
| 0.00247362 | 5 |
| 0.0049865 | 6 |
| 0.0100123 | 7 |
| 0.0200638 | 8 |
| 0.0401668 | 9 |
| 0.0803729 | 10 |
| 0.160785 | 11 |
| 0.32161 | 12 |
| 0.643258 | 13 |
| 1.28656 | 14 |

**Figure S6.** The time of lipid self-assembly as a function of doubling steps, where 14 doubling steps (the required number of steps to form an eLNP with a 40 nm diameter and 3 nm bilayer) equates 1.29 ms.

*Organic strip width calculation*

Using the initial feed composition of lipids (C_lipids, feed_ = 12 mg/mL) and calculating the mass of lipids that exist in the feed, the volume of a vesicle-forming region containing the ensemble of lipids needed to make an eLNP of 40 nm diameter and 3 nm bilayer thickness can be calculated. To start, the mass of lipids (m_lipids_) that exist in the feed is calculated using **Eqn. E9** from the turbulent flow *in situ* loading calculations. **Eqn. E9** can then be used to calculate the assembly sphere volume (V_vesicle-forming region_) in nm^3^ of lipids needed to make an eLNP:

$$V{}_{vesicle-forming region}=\frac{m{}_{Lipids}}{C_{lipids, feed}}\times{10}^{21}$$

(E9)

Assuming that the volume is spherical, the radius (R_vesicle-forming region_ [=] nm) can be backed out using the volume of a sphere in **Eqn. E10**:

$$R_{vesicle-forming region}= \sqrt[3]{\frac{3}{4\pi}V_{vesicle-forming region}}$$

(E10)

For the given starting parameters, this results in a radius of 63.3 nm or a 126.6 nm diameter vesicle-forming region required to make an eLNP. This (L =127 nm = 1.27 x 10^-7^ m) will act as the thickness of the strips analyzed in the following calculations.

*RNA diffusion from the aqueous to the organic channel*

To start, the diffusivities of trace ethanol in water (1.2 x 10^-5^ cm^2^/s) and trace water in ethanol (1.0 x 10^-5^ cm^2^/s) are averaged (D_avg_ = 1.1 x 10^-9^ m^2^/s)^1^. This is done to determine how long it takes water and ethanol to mix. Using **Eqn. E11**, the time for ethanol and water to fully mix is calculated.

$$\tau=\frac{L^{2}}{D}$$

(E11)

For a strip width of 127 nm, the time for ethanol and water to mix is determined to be 15 µs. Due to over 50x difference between the time for ethanol and water to mix and lipid self-assembly, ethanol-water diffusivity is considered negligible.

Calculating the diffusivity of RNA of molecular weights ranging from 3.4 to 2000 kDa using **Eqn. E7**, and using the lipid self-assembly time in **Eqn. E11**, the length in which RNA diffuses (L_RNA_) can be assessed. Starting halfway through the strip, the distance an RNA can travel is compared to the overall width of the aqueous channel (W­­_aqueous_ = 0.075 mm) to determine that RNAs in what percentage of the aqueous channel (P_Aqueous_) (i.e. what percentage of total RNAs) could diffuse halfway through the strip of lipids. This is seen in **Eqn. E12**:

$$P_{Aqueous}=\frac{\left( L_{RNA}-\frac{L}{2} \right)}{W_{Aqueous}}\times100\%$$

(E12)

Doing this analysis for different molecular weights of RNA, where L_RNA_ changes due to a change in RNA diffusivity, **Figure S7** shows how for all molecular weights assessed, that less than 1% of the RNA residing in the aqueous channel will likely undergo *in situ* loading (ISL).

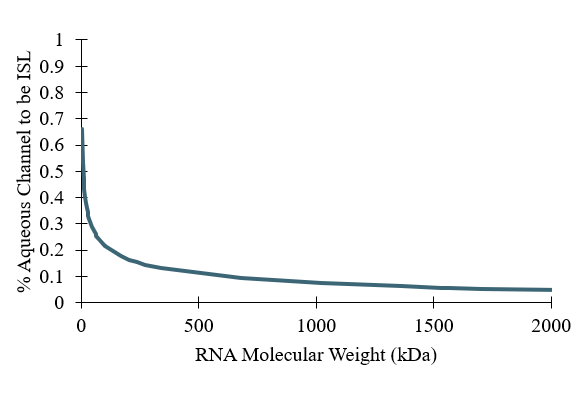

**Figure S7.** Percentage of the RNA within the aqueous channel able to diffuse halfway through one strip of lipids during the time it takes an eLNP to form as a function of RNA molecular weight (kDa). Less than 1% of all molecular weight RNAs will undergo *in situ* nanoprecipitation with lipids. The remaining RNA will likely be *post hoc* loaded into eLNPs through RNA bridged fusion events.

*Primary vesicle fusion number calculation*

The liposome-like primary vesicles formed at low pH (40nm, 3.5nm bilayer) have the same outer surface area as a filled 40nm sphere, but only ~43% as much mass (see above for the calculation of the bilayer volume; this volume is 43% as much as a filled sphere of the same outer radius). Assuming a neutralized LNP post-fusion has a diameter of 80nm, the number of primary vesicles required to fuse to obtain this mass is given by $N=\frac{m_{80nm LNP}}{{0.57m}_{40nm filled sphere}}=\frac{{(40nm)}^{3}}{0.57{(20nm)}^{3}}=$ ~14. If a mixed population of primary vesicles has e.g. 1 out of every 14 vesicles loaded with a high MW mRNA, the likelihood that upon fusion, each loaded vesicle will fuse with exactly 13 empty vesicles to form a homogeneous population of complete LNPs is vanishingly small.

**Reference:**

1. Khattab, I.S., Bandarkar, F., Fakhree, M.A.A. et al. Density, viscosity, and surface tension of water+ethanol mixtures from 293 to 323K. *Korean J. Chem. Eng.* 29, 812–817 (2012). https://doi.org/10.1007/s11814-011-0239-6
